# Batf-mediated Epigenetic Control of Effector CD8+ T Cell Differentiation

**DOI:** 10.1101/2021.01.04.425241

**Authors:** Hsiao-Wei Tsao, James Kaminski, Makoto Kurachi, R. Anthony Barnitz, Michael A. DiIorio, Martin W. LaFleur, Wataru Ise, Tomohiko Kurosaki, E. John Wherry, W. Nicholas Haining, Nir Yosef

**Affiliations:** Department of Pediatric Oncology, Dana-Farber Cancer Institute, Boston, MA, USA; Broad Institute of MIT and Harvard, Cambridge, MA, USA; Center for Computational Biology, University of California, Berkeley, Berkeley, CA, USA; Department of Molecular Genetics, Graduate School of Medical Sciences, Kanazawa University, Kanazawa, Japan; Evergrande Center for Immunologic Diseases, Harvard Medical School and Brigham and Women’s Hospital, Boston, MA, USA; Division of Medical Sciences, Harvard Medical School, Boston, MA, USA; Department of Microbiology and Immunobiology, Harvard Medical School, Boston, MA, USA; Department of Microbiology and Institute for Immunology, University of Pennsylvania Perelman School Medicine, Philadelphia, PA, USA; Ragon Institute of Massachusetts General Hospital, Massachusetts Institute of Technology, and Harvard University, Boston, MA, USA; Department of Electrical Engineering and Computer Science, University of California, Berkeley, Berkeley, CA, USA; Division of Pediatric Hematology and Oncology, Children’s Hospital, Boston, MA, USA; Chan Zuckerberg Biohub, San Francisco, CA, USA; Laboratory of Lymphocyte Differentiation, WPI Immunology Frontier Research Center, Osaka University, Osaka, Japan; Laboratory for Lymphocyte Differentiation, RIKEN Center for Integrative Medical Sciences, Kanagawa, Japan

## Abstract

The response of cytotoxic T cells to their cognate antigen involves rapid and broad changes in gene expression that are coupled with extensive chromatin remodeling. Here, we study the mechanisms by which the basic leucine zipper ATF-like transcription factor Batf helps regulate this process. Through genome-scale profiling, we observe critical roles for Batf in inducing transcriptional changes in stimulated naive cells, while affecting the chromatin at several levels, namely binding of key transcription factors, accessibility, and long range contacts. We identify a critical network of transcription factors that cooperate with Batf, including its binding partner Irf4, as well as Runx3 and T-bet, and demonstrate its synergistic activity in initiating aspects of the effector T cells’ transcriptional and epigenetic program in ectopically-induced fibroblasts. Our results provide a comprehensive resource for studying the epigenomic and transcriptomic landscape of effector differentiation of cytotoxic T cells.

## INTRODUCTION

When CD8+ T cells encounter their cognate antigens, they embark on a process of rapid clonal expansion and differentiation, generating a large population of effector cells characterized by marked changes in gene expression. The transcriptional changes that occur during effector differentiation are associated with massive shifts in chromatin accessibility, reflecting large-scale changes in both regulation of and transcription from gene loci (Kaech and Cui, 2012; Sen et al., 2016; Wong and Pamer, 2003).

The basic leucine zipper activating transcription factor (ATF)-like transcription factor (Batf) is a well-characterized transcription factor (TF) involved in the development, differentiation and function of many immune cell subsets, including dendritic cells (DC), B cells, T helper 2 (Th2), Th9, Th17, follicular T helper (Tfh) and CD8+ T cells (Betz et al., 2010; Ciofani et al., 2012; Ise et al., 2011; Jabeen et al., 2013; Kurachi et al., 2014; Kuwahara et al., 2016; Sahoo et al., 2015; Schraml et al., 2009). Batf is reported to be induced by various stimuli in T cells including TCR stimulation and cytokine signaling, and contributes to T cell function through divergent mechanisms (Kuroda et al., 2011; Man et al., 2017; Pham et al., 2019; Xin et al., 2015). In Tfh, Batf is able to recruit Ctcf in an Ets1 dependent manner to establish the chromatin landscape for initiating the earliest phase of differentiation. This mechanism is independent of interferon regulatory factor 4 (Irf4) (Pham et al., 2019). However, by cooperating with Irf4, Batf is critical for initiating the differentiation of Th17 cells, acting as so-called pioneer factors (Ciofani et al., 2012; Li et al., 2012; Murphy et al., 2013), which regulate chromatin remodeling and facilitate the binding of other factors (Zaret and Carroll, 2011). Batf and Irf4 are capable of binding a composite AP-1-IRF DNA element (AICE) binding sites, which served to explain their interdependence (Glasmacher et al., 2012; Li et al., 2012). Like Batf knockout (KO) mice, mice deficient for Irf4 exhibit defects in Tfh cell differentiation, Th17 differentiation, CD8+ T cell response to LCMV infection, and B cell class switch recombination (Bollig et al., 2012; Brüstle et al., 2007; Grusdat et al., 2014; Klein et al., 2006; Mudter et al., 2011; Sciammas et al., 2006).

Previously, we demonstrated that Batf is also necessary for the earliest differentiation and expansion of CD8+ T cells when they encounter an antigen *in vivo* (Kurachi et al., 2014). Knockout of *Batf* in CD8+ T cells results in proliferation defects, changes in mRNA expression of other transcription factors and abnormal cytokine production. Furthermore, chromatin immunoprecipitation followed by sequencing (ChIP-seq) of Batf in *in vitro* differentiated effector CD8+ T cells suggested that it may colocalize with other transcription factors and possibly facilitate their binding. Although Batf has been shown to be able to initiate the CD4+ T cell chromatin reorganization and differentiation in the Tfh-prone environment (Pham et al., 2019), how Batf establishes the cell identity and effector function in CD8+ background has not been described in a systematic way. Furthermore, the mechanisms by which Batf coordinates the epigenomic and transcriptomic events during CD8+ T cell activation and differentiation, how Batf synchronizes with other transcription factors, and the extent to which Irf4 is required for these aspects of BATF function, are still unclear.

In order to explore these questions, we utilized multiple types of genome-scale assays to map the effects of Batf deficiency on the transcriptome and epigenome of differentiating and effector CD8+ T cells. We first evaluated the relevant transcriptional changes using RNA-seq, comparing Batf deficient to normal cells. We then investigated the respective chromatin-level changes using assay for transposase accessible chromatin sequencing (ATAC-seq) and ChIP-seq of a set of critical TFs to produce a dataset of 96,490 genomic regions with accessible chromatin or bound TFs, and identified long-range contacts among these regions using Hi-C chromatin immunoprecipitation (HiChIP). Through systematic analysis of these transcriptional and epigenetic datasets, we identified a Batf-dependent transcriptional program that establishes effector differentiation. This program consisted of major changes in chromatin accessibility and three dimensional structure, binding of key TFs, and corresponding changes in gene expression. We identified a subset of key TFs that are co-localized and affected by Batf in T cells (Irf4, Runx3, and T-bet). We then demonstrated that ectopic expression of these TFs along with Batf in fibroblasts, but not Batf alone, is sufficient to reproduce aspects of the epigenetic (chromatin accessibility and loops) and transcriptional characteristics of CD8+ effector T cells.

Taken together, our results not only map the binding of critical TFs but also provide a comprehensive resource for studying the epigenome of differentiating CD8+ T cells. They also yield an in-depth view of the manner by which Batf plays its role in directing effector CD8+ T cell differentiation after engagement with antigen, ultimately reaching proper cytotoxic T cell response against pathogens.

## RESULTS

### Loss of Batf has broad effects on gene expression and chromatin accessibility during CD8+ T cell differentiation

To understand the role of Batf in regulating the differentiation of effector CD8+ T cells, we generated inducible Batf conditional knockout (cKO) CD8+ T cells derived from the P14 TCR transgenic mouse, in which the CD8+ T cells recognize the GP_33-41_ lymphocytic choriomeningitis virus (LCMV) epitope (Pircher et al., 1987). The Batf cKO P14 CD8+ T cells generated in donor mice were adoptively transferred to recipient mice one day before infection with the Armstrong (Arm) strain of mouse LCMV, which causes acute infection (Figures **<u>1A</u>** and **<u>S1A</u>**). Wild type (WT) P14 CD8+ T cells were also sorted and co-transferred to recipient mice as control.

**Figure 1.**
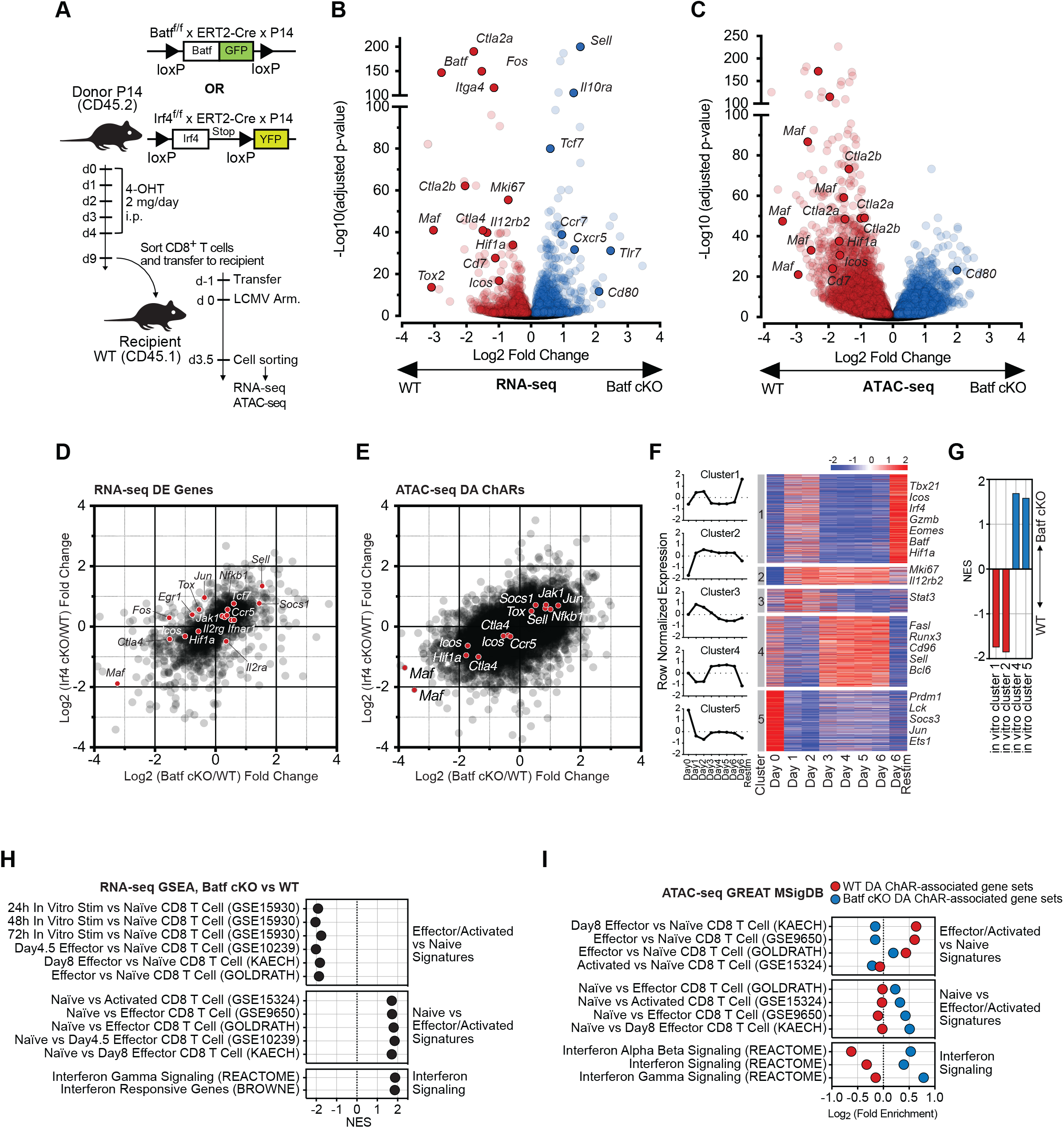
Batf and Irf4 remodel chromatin accessibility at key regulatory regions during effector CD8+ T differentiation. **(A)** Schematic of Cre-lox conditional KO system and LCMV acute infection mouse model. **(B)** Volcano plot of log2 fold change in gene expression (Batf cKO/WT) and adjusted p-values for differential expression tests in Batf cKO CD8+ T cells. A subset of genes with important T cell functions are labeled. **(C)** Volcano plot of fold change in chromatin accessibility in a chromatin ChAR, measured by Tn5 cuts in locus (Batf cKO/WT), and adjusted p-values for tests of differential accessibility. ChARs were obtained by calling ATAC-seq peaks in the WT or Batf cKO conditions with MACS2. ChARs were mapped to putative target genes with GREAT, and a subset of ChARs mapped to genes with important T cell functions are labeled. **(D)** Scatterplot showing correlation between Batf cKO DE gene fold-change and Irf4 cKO DE gene fold-change. The universe of genes is limited to those with a significant change in expression (adjusted p-value < 0.05) in one of the two experiments. **(E)** Scatterplot showing the correlation between fold-change of Batf cKO and Irf4 cKO DA ChARs. The universe of genes is limited to those with a significant change in expression (adjusted p-value < 0.05) in one of the two experiments. A subset of ChARs mapped to putative target genes with important T cell functions are highlighted. **(F)** Transcriptomic profiling of *in vitro* cultured CD8+ T cells. Gene expression was measured with RNA-seq daily, for six days. Row normalized expression counts are shown pre-stimulation (day 0), after primary anti-CD3/CD28 stimulation (days 1 to 6), and after PMA/ionomycin restimulation (day 6 restimulated). Genes were clustered into 5 groups using *hierarchical* clustering. Average expression trajectory for each cluster is shown on the left. **(G)** Normalized enrichment score (NES) for GSEA in the clusters shown in **(F)** in Batf cKO vs WT conditions. **(H)** NES for selected immunologic signatures of Batf cKO vs WT DE gene sets. Higher NES indicates enrichment in Batf cKO. Only results with adjusted p-value < 0.05 are shown. **(I)** Fold enrichment for selected immunologic signatures of WT vs Batf cKO DA ChARs analyzed using GSEA C7 immunological signature gene sets and GREAT. WT DA ChAR-associated gene sets are shown in red and Batf cKO DA ChAR-associated gene sets are shown in blue.

Our previous study demonstrated that Batf KO P14 TCR transgenic CD8+ T cells cannot control acute infection of LCMV, and instead exhibited cell death four days post infection (Kurachi et al., 2014). To investigate viable CD8+ T cells in this model, we therefore harvested the Batf cKO cells at 3.5 days post infection, shortly before the anticipated population collapse, and performed RNA-seq and ATAC-seq in order to compare the Batf cKO to the WT control. RNA-seq showed that deletion of Batf resulted in significant transcriptional changes, with 2,552 genes affected (1,292 increased and 1,260 decreased; adjusted p-value < 0.05; see **Methods** and Figure **<u>1B</u>**, <u>Table **1**</u>). Batf cKO CD8+ T cells had a marked decrease in mRNA levels of TFs and signaling molecules that are important for T cell activation and function, including *Fos*, *Ctla4*, *Maf*, *Il12rb2*, *Hif1a* and *Cd7*. Genes with increased expression in the cKO samples included ones that are characteristic of naive T cells such as *Sell*, *Tcf7*, and *Ccr7* as well as other genes involved in T cell function such as *Il10ra* and *Cxcr5*.

To determine if loss-of-function of Batf also affected the chromatin landscape in those antigen-exposed CD8+ T cells, we investigated the changes in chromatin accessibility observed between the WT and the cKO conditions using ATAC-seq. We first used peak calling to map all loci that were sufficiently accessible in at least one condition (Figure **<u>S1A</u>** and **Methods**), identifying a total of 38,055 chromatin accessible regions (ChARs). We then searched for ChARs that were differentially accessible (DA) (see **Methods,** <u>Table **2**</u>). Overall, 6,011 ChARs were more accessible in the Batf cKO condition, while 6,605 ChARs were less accessible (Figure **<u>1C</u>**). These results showed that Batf is required to establish both the chromatin landscape and transcriptional program of activated CD8+ T cells. Unlike CD4+ T cells (Pham et al., 2019), the changes in chromatin accessibility and transcriptome in Batf-deficient CD8+ T cells were bidirectional which indicated dual roles of Batf as a possibly indirect positive or negative regulator.

We further confirmed the relationship between chromatin changes and gene expression by associating each DA ChAR with nearby genes using GREAT (<u>Table **3**</u> and **Methods**). We found that a significant proportion (58.7%) of the differentially expressed (DE) genes were associated with at least one DA ChAR (48.5% if we only associate a ChAR with its nearest gene). This level of overlap was higher than expected by chance (fold enrichment = 1.24, p < 0.001) (Figure **<u>S1B</u>**). We also observed a limited quantitative correspondence between the amount of change in mRNA expression and change in accessibility around each gene locus (Pearson correlation of 0.187 (p < 1e-4) if we consider all loci, and of 0.622 (p < 1e-7) if we only consider ChARs that resided in the promoter of their putative target gene Figure **<u>S1C</u>**). The set of genes that were downregulated in Batf cKO and associated with DA ChARs with reduced accessibility included several key TFs and signaling molecules in effector T cells, such as *Maf, Ctla4, Ctla2a, Ctla2b, Hif1a, Icos and Cd7.* Conversely, ChARs within and around the promoter of the *Sell* gene, which is highly expressed by naive cells and encodes for the L-selectin CD62L tended to be more accessible in Batf cKO. Consistently, we observed a three-fold increase of mRNA expression of that gene in the cKO condition (Figure **<u>S1D</u>**).

Taken together, these results point to a coordinated transcriptome- and epigenome-wide dependence of CD8+ T cells differentiating *in vivo* on Batf. This is consistent with previous findings that proper function of this factor is critical for mounting an effective anti-viral or other adaptive immune responses (Grusdat et al., 2014; Xin et al., 2015).

### The effects of Irf4 on the transcriptome and epigenome overlap with those of Batf

The TF Irf4 is a well characterized binding partner of Batf (Li et al., 2012), and the importance of the interaction between these factors has been demonstrated in several processes, including ones in T helper cells (Ciofani et al., 2012; Li et al., 2012), tissue-resident regulatory T cells (Vasanthakumar et al., 2015) and cytotoxic T cells (Grusdat et al., 2014). To study the role of Irf4 in shaping the transcriptome and chromatin landscape of activated CD8+ T cells, and to investigate whether it affects a similar set of genes and loci as Batf, we generated Irf4 cKO P14 T cells *in vivo*, using the same mouse LCMV model and experiment timing described for Batf (Figures **<u>1A</u>** and **<u>S1A</u>**), and conducted RNA-seq and ATAC-seq (Figures **<u>S1E</u>** and **<u>S1F</u>,** Tables **1** and **2**).

We found that the transcriptional effects of Irf4 loss were largely consistent with that of Batf (Pearson correlation of 0.506 between the induced fold changes; p < 1e-4; Figure **<u>1D</u>**). For example, *Sell*, *Socs1*, *Nfkb1* and *Tcf7* were similarly up-regulated in both Batf cKO and Irf4 cKO whereas *Ctla4* and *Maf* were down-regulated in both cKO cells. The effects on chromatin accessibility were largely consistent as well (r = 0.494; p< 1e-4; Figure **<u>1E</u>**). The combined effects in both modalities was observed in several key loci, such as *Ctla4* and *Maf*, which exhibited marked decrease in accessibility and mRNA levels in both perturbations, while *Sell, Socs1* and *Nfkb1* showed a marked increase in both modalities and both perturbations. It should be noted, however, that generally the loss of Batf led to an effect of stronger magnitude on chromatin accessibility compared to Irf4 cKO while they had comparable affected gene numbers (Figures **<u>S1G, S1H</u>** and **<u>S1I</u>**). This, along with the partial overlap between the effects (e.g., on the expression of the TFs Jun and Fos, Figure **<u>1D</u>**), suggests that although Batf and Irf4 shared many aspects of chromatin and transcriptional changes when knocking out, they may also have distinct contributions to the epigenetic and transcriptional rewiring required for T cell activation.

### The Batf dependent transcriptome and epigenome are associated with effector T cell differentiation

To better connect between the observed effects of Batf to the process of T cell response to antigens, we produced, as a reference, a time course gene expression dataset from *in vitro* CD8+ culture, from naive to effector. To this end, naive CD8+ T cells were isolated from WT mice, stimulated with anti-CD3/CD28, and assyed up to 6 days post stimulation with an additional restimulated sample at day 6 (using PMA/ionomycin; see **Methods**). We first identified genes whose expression changed over time and divided these genes into five prototypical clusters of response (Figure **<u>1F</u>** and **Methods**). Using GSEA (<u>Table **3**</u>), we then found that the set of genes that were up-regulated in Batf cKO significantly overlapped with clusters of genes that had reduced expression after stimulation (Clusters 4 and 5, adjusted p < 0.01), including *Fasl* (apoptosis-inducing gene (Strasser et al., 2009)), *Scos3* (cytokine signaling regulator (Brender et al., 2007)) and *Sell* (naive T cell marker (Arbonés et al., 1994)). Conversely, genes that were down-regulated in the absence of Batf were associated with clusters that become induced post primary stimulation (Figure **<u>1G</u>**, Clusters 1 and 2, adjusted p < 0.01), including the costimulatory receptor *Icos (Dong et al., 2001),* CD8+ effector response genes such as *Hif1a* (Phan and Goldrath, 2015), cytokine receptors that are related to effector generation such as *Il12rb2* (Danilo et al., 2018) and the cell proliferation marker *Mki67* (Gerdes et al., 1984). From the epigenome perspective, we found that regions which were more accessible in the Batf cKO were most strongly associated with clusters of reduced expression after stimulation (using GREAT; Clusters 4 and 5, q value < 1e-5; <u>Table **4**</u>). Regions that were more accessible in the WT had a less clear trend, with significant overlap with cluster 1 (induced expression post stimulation) and cluster 4 (decreased; q value < 0.05). Results for Irf4 cKO were similar (Figure **<u>S1J</u>** and <u>Table **4**</u>), suggesting that deficiency of Batf or Irf4 impairs the expression of TCR-inducible genes and the subsequent transition from naive to effector state. These results accord with our previous work whereby loss of Batf led to defects in the differentiation and expansion of stimulated CD8+ T cells *in vivo (Kurachi et al., 2014).*

To further explore this hypothesis, we utilized a compendium of published data of transcriptional changes in CD8+ T cells under different conditions. We tested whether these previously observed changes overlap with the effects we observed of Batf on the transcriptome and epigenome (using GSAE and GREAT, see **Method**; Figures **<u>1H</u>**, **<u>1I</u>**, **<u>S1J</u>** and **<u>S1K</u>**, Tables **<u>3</u>** and **<u>4</u>)**. Both data modalities clearly associated the effects of Batf in our data with previously observed differences between naive and effector CD8+ T cells. For instance, genes that were down-regulated in the absence of Batf in our data tend to be up-regulated in the effector state. Similarly, genes that were proximal to ChARs that lost accessibility in Batf cKO tended to be over-expressed by the effector and activated populations. Results from Irf4 cKO also showed a similar trend towards the naive CD8+ state when comparing to the effector state (Figure **<u>S1J</u>** and **<u>S1K</u>**). Using a similar GSEA procedure we also found that the sets of Batf-affected genes and loci significantly overlap with genes and loci that have been associated with interferon signaling and response. These results are concordant with our previous work showing abnormal interferon gamma (IFNg) production in Batf KO cells *in vivo* (Kurachi et al., 2014). Interestingly, Irf4 cKO did not display a strong impact on interferon signalling pathways at transcriptomic level (Figure **<u>S1J</u>**), suggesting a potential mechanism of Batf that is independent of Irf4. Finally, analysis of DNA binding motifs in Batf-dependent ChARs revealed a significant over-representation of motifs that are associated with transcription factors or complexes that are important for (but not necessarily specific to) T cell function such as the tryptophan cluster (IRF, ETS and etc.), high mobility group (HMG), and homeodomain. We also observed strong enrichment of basic leucine zipper (bZIP) motifs, which can be potentially bound by Batf itself (Figure **<u>S1L,</u>** <u>Table **4**</u>).

Overall, these results serve to place the effects of Batf and Irf4 in the context of the process of CD8+ T cell effector differentiation. We observed, in both cases, that the transcriptional and epigenetic changes induced by the loss of Batf or Irf4 had a significant level of similarity to the ones observed prior to the antigen exposure and activation of naive T cells. These results therefore suggest that in the cKO conditions, the cells were prone to stall in a naive state compared to WT.

### *In vitro* profiling of transcription factors reveals genome-wide dependence on Batf

While our motif analysis suggested plausible dependence of other transcription factors on Batf function, it affords a limited view of the actual binding landscape (Yue et al., 2014). For a more direct analysis we used ChIP-seq of WT and Batf KO P14 CD8+ T cells (Kurachi et al., 2014) cultured *in vitro* to identify key binding locations and evaluate their dependence on Batf. To this end, naive cells were stimulated with anti-CD3/CD28, cultured for 6 days and restimulated with PMA/ionomycin (similarly to Figure **1F**; see **Methods**). We assayed a list of transcription factors whose presence is important for T cell function and development, including the CD8+ lineage determinant, T-bet (Intlekofer et al., 2005; Sullivan et al., 2003), the factor for memory generation, Eomes (Intlekofer et al., 2005; Pearce et al., 2003), factors critical for CD8+ development such as Runx3 and (Cruz-Guilloty et al., 2009; Shan et al., 2017), Ets1 (Grenningloh et al., 2011; Zamisch et al., 2009), a factor required for maintaining effector function, Fosl2 (Lingel et al., 2017; Lund et al., 2007; Stelekati et al., 2018), the Batf binding partner and transactivator Jund (Li et al., 2012), and the cytokine signaling transducers such as Stat3 (Cui et al., 2011; Kujawski et al., 2010) and Stat5 (Tripathi et al., 2010), in addition to Batf and Irf4. The set of putative binding locations (peaks) in each sample was identified using MACS2 (Zhang et al., 2008) (<u>Table **2**</u>, Figures **<u>2A</u>** and **<u>S2A</u>**).

**Figure 2.**
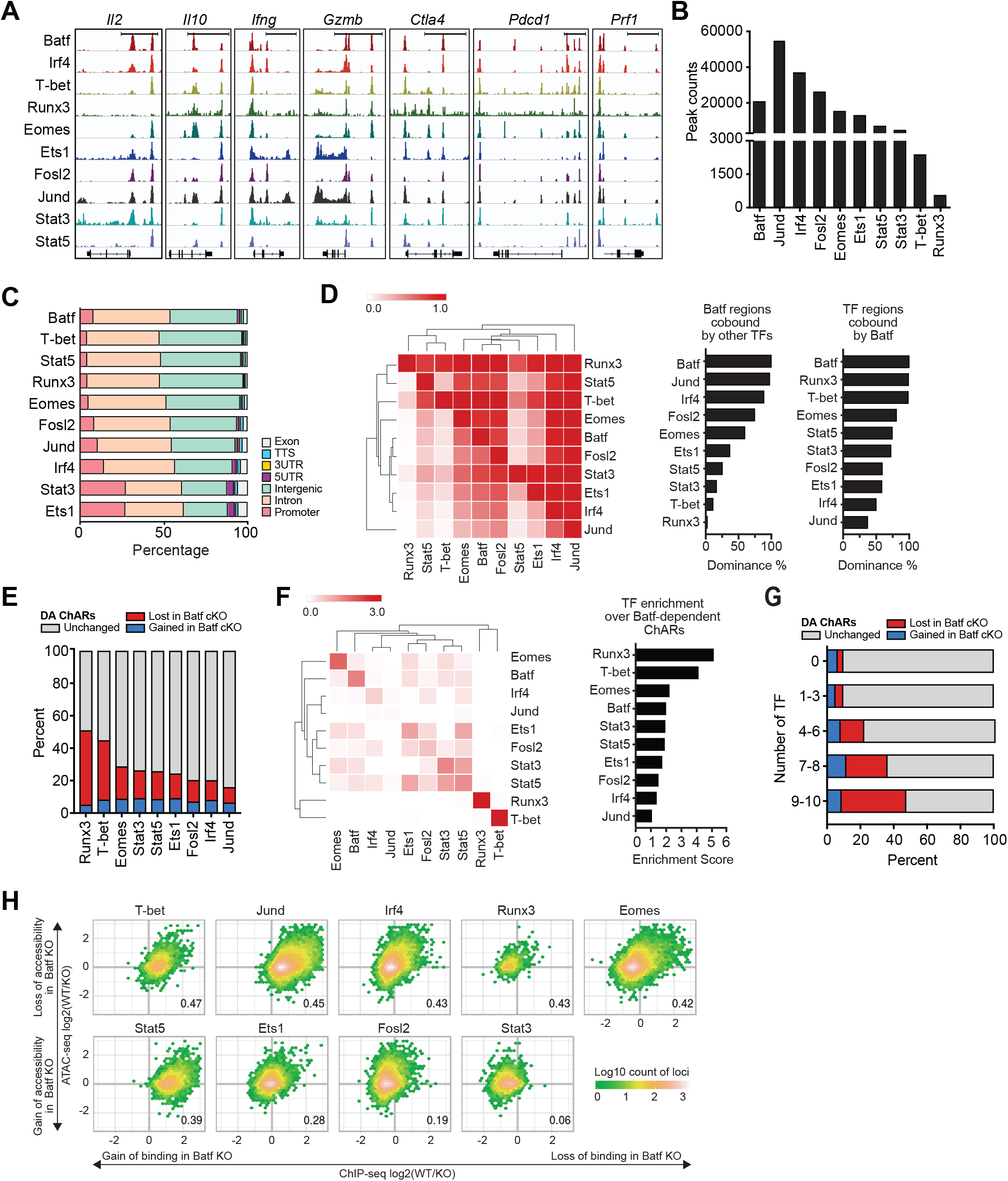
Batf is required for the binding of key transcription factors in CD8 T cells and contributes to CD8 T cell phenotypes by cooperating with Runx3 and T-bet. **(A)** Representative pileup of ChIP-seq fragments for main TFs in WT CD8+ T cells around selected gene loci via IGV. Scale bars represent 5 kb. **(B)** The total counts of peaks for each TF in *in vitro* WT CD8+ T cell samples. **(C)** Percentage of TF peaks across genomic features for WT samples. **(D)**(Left heatmap) Percentage of each row TF’s peaks that are co-bound by the column TF. (Center barplot) Percentage of each TF’s peaks that are co-bound by BATF. (Right barplot) Percentage of Batf peaks that are co-bound by each TF. **(E)** TF ChIP-seq peaks stratified by change in accessibility in Batf cKO ATAC-seq: lost, gained, or unchanged. **(F)** Fold enrichment of TF binding over the DA ChARs lost in Batf cKO samples. Off-diagonal entries in the heatmap show the enrichment of regions co-bound by the two respective TFs. Diagonal entries in the heatmap and the barplot to the right show the enrichment for individual TFs. See **Method** for details of fold enrichment calculation. **(G)** Percentage of TF-bound regions, categorized by number of TFs bound, each stratified by change in accessibility in Batf cKO ATAC-seq: lost, gained, or unchanged. **(H)** Plot of relationship between change in TF binding (x-axis) and change in DNA accessibility (y-axis) for main TFs at loci bound by the TFs in WT or Batf KO conditions. The log2(WT/Batf KO) values for binding and accessibility were estimated by applying csaw and DESeq2 to ChIP-seq and ATAC-seq data, respectively, and these points are binned into hexes colored by density of loci falling within them. Only loci exhibiting significant change in either binding or accessibility are shown (adjusted p-value< 0.05). TFs are arranged in order of decreasing Pearson correlation, which is shown on the lower-right of each plot.

Starting with the WT samples, we found a broad range in the extent of binding, from widespread (tens of thousands of loci for Jund and Irf4), to more specific (few thousand loci or less for Runx3 and T-bet; Figure **<u>2B</u>**), with most sites residing in non-coding regions (Figure **<u>2C</u>**, <u>Table **2**</u>). We also observed a large extent of colocalization between the TFs in our dataset (Figure **<u>2D</u>**). For instance, and as expected, Batf bound loci significantly overlapped with those of its well-characterized partners, Irf4 and Jund (Figure **<u>2D</u>** middle). We also observed that most of the binding regions of Runx3 and T-bet were proximal to a Batf binding site (Figure **<u>2D</u>** right). Furthermore, most TFs exhibited a clear preference for localizing near ChARs that lost accessibility in Batf KO (more than expected by chance, p < 0.001; Figure **<u>2E</u>** and **Methods**). This effect was most pronounced in regions bound by Runx3 and T-bet (Figure **<u>2F</u>**) or in “hot” regions occupied by multiple TFs (Figure **<u>2G</u>**, <u>Table **2**</u>). A similar trend was observed with DA ChARs in the Irf4 cKO (Figure **<u>S2B</u>**). Taken together, these results demonstrate that a group of TFs that are critical for CD8+ differentiation and function tend to operate in overlapping regions of the genome, and that Batf and Irf4 have a broad effect on the accessibility of these regions. We next investigated whether the affinity of binding in those regions indeed changed as a consequence of Batf deficiency.

To explore this we plotted, for each TF, the changes in its binding signal versus the changes in accessibility in all loci where it bound next to Batf and where at least one of the two signals (evaluated by ATAC-seq and ChIP-seq) changed significantly (Batf KO vs. WT; adjusted p-value < 0.05; Figure **<u>2H</u>**, <u>Table **2**</u>, and **Methods**). The majority of TFs had a significant positive association between changes in accessibility and changes in binding, suggesting that a nearby Batf binding event may help facilitate, directly or indirectly, their own binding by means of chromatin organization. The most pronounced loss of binding was observed with Jund, which is capable of dimerizing and co-binding with Batf (Murphy et al., 2013), and with Stat5, which mediates IL-2 signaling and consequently differentiation and homeostasis (Ross and Cantrell, 2018). Interestingly, and in contrast to Stat5, the binding of Stat3 has primarily increased upon loss of Batf function, where a substantial part (48.3%) of the gained sites overlapped with a loss in Stat5 binding (using csaw for differential binding with FDR < 5%; See <u>Table **3**</u> and **Methods**). An example for such Stat5 to Stat3 switch was found proximal to the BCL-2 family member Bim (*Bcl2l11*), which is regulated in T cells by JAK/STAT signaling (Bouillet et al., 2002) and whose expression increased in Batf cKO (adjusted p-value < 0.001; Figure **S6A**). These results accord with previous reports on competitive binding and opposing regulatory effects of these two TFs (Walker et al., 2013) and suggest a role for Batf in maintaining their opposing effects in activated CD8+ T cells. Unlike most other TFs we profiled, the changes in the binding of Stat3 in the absence of Batf did not have a strong association with changes to accessibility. This may suggest other means in which Batf exerts its effects in Stat3 sites, such as changes in the activity of binding competitors (Sadreev et al., 2018; Walker et al., 2013; Wingelhofer et al., 2018) and/or in post translational modifications at nearby histone tails (Chan and Maze, 2020). Notably, we observed a similar trend of low correlation between changes in binding versus changes in accessibility in the case of Fosl2. This result accords with previous reports on antagonistic relationship between Fosl2 and Batf in Th0 and Th17 conditions, whereby Fosl2 binding sites are overlapped with those bound by Batf, thus competing for the same AP-1 sites (Ciofani et al., 2012).

Taken together, our results suggest that Batf may, at least partially, exert its effects on CD8+ effector differentiation by modulating the binding landscape of critical TFs and may be dependent or independent of chromatin accessibility. Of note, out of the ten TFs we studied, Runx3 and T-bet had the largest overlap with Batf binding sites and with Batf-dependent DA ChARs, while showing clear positive association between these two signals.

### Batf-mediated chromatin loops are associated with Batf-dependent accessible regions and transcription factor binding

While we investigated the effects of Batf on the chromatin by mapping local changes in accessibility and TF binding (Figure **1** and Figure **2**), these effects may also be associated with higher-order alterations, namely long-range contacts between otherwise distant genomic regions (Pham et al., 2019). We therefore used HiChIP (Mumbach et al., 2016) for the *in vitro* differentiated effector CD8+ T cells (naive CD8+ T cells stimulated with anti-CD3/28, cultured for 6 days and restimulated with PMA/ionomycin, same condition as in figure **1F**, figure **2**, see **Methods**) to identify chromatin loops that are likely dependent on the presence of Batf. For the purpose of comparison, we used the CCCTC-binding factor (Ctcf), which plays a housekeeping role in defining genomic boundaries in many types of mammalian cells, including T cells (Rowley and Corces, 2018). Starting with a global perspective, we observed a significant level of overlap between regions nearby Batf-dependent loops and binding locations of TFs in our study. We also found a significant overlap between Batf-dependent loops and ChARs that lost accessibility in the absence of Batf or Irf4. Notably, the observed overlaps were generally more pronounced than the ones observed with Ctcf-anchored loops (Figure **<u>3A</u>**, <u>Table **5**</u>).

**Figure 3.**
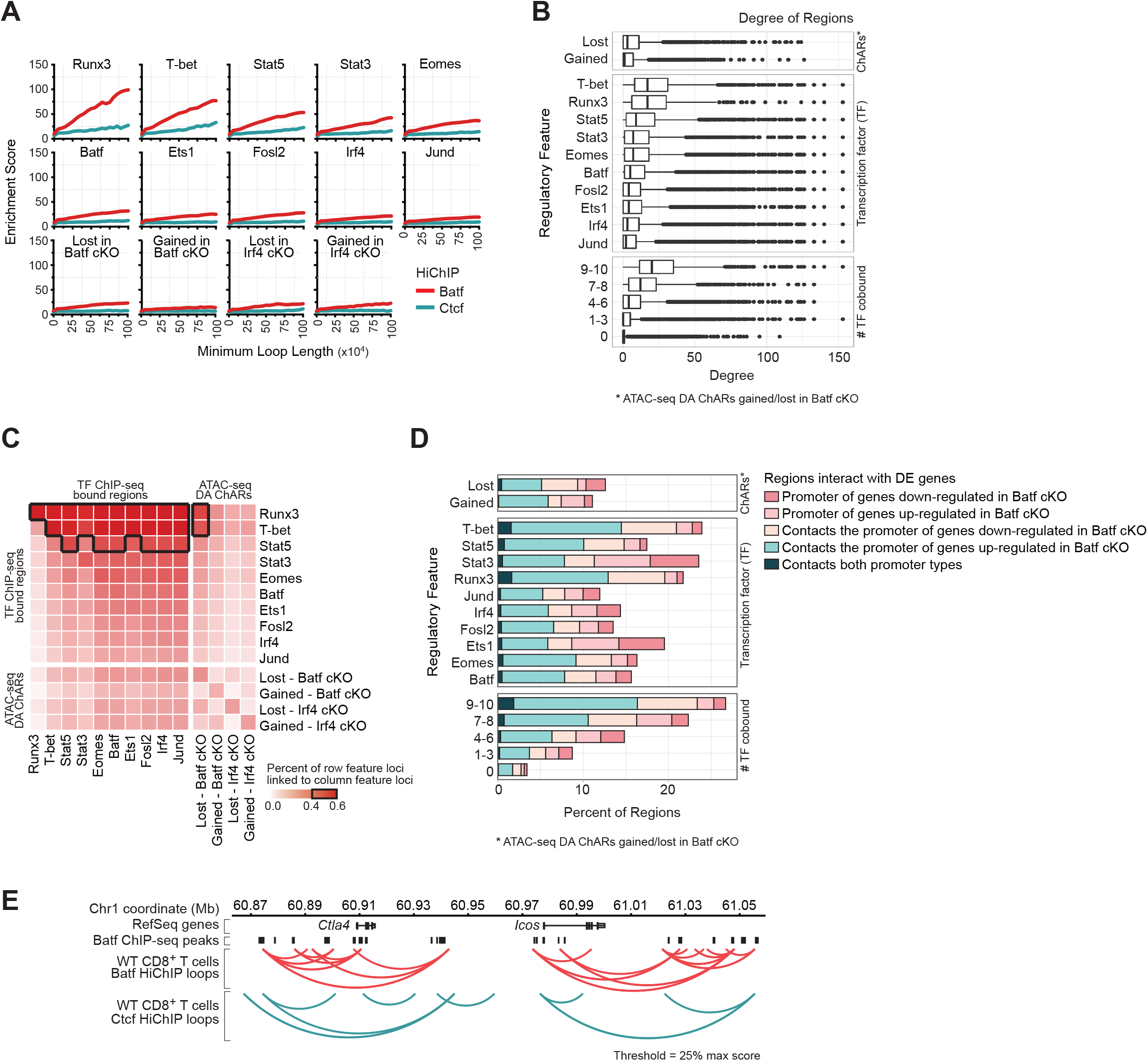
Differential accessible ChARs and loci bound by key transcription factors are linked by Batf loops in CD8 T cells. **(A)** Fold enrichment of area spanned by HiChIP loop anchors for Batf (red) and Ctcf (blue-green) in area spanned by key CD8+ TFs and ChARs gained and lost in Batf cKO and Irf4 cKO CD8+ T cells. The enrichment score is defined as “percent of genome spanned by feature / percent of HiChIP loop anchor area spanned by feature”. **(B)** Boxplot of number of Batf-associated contacts to other regions, categorized by epigenetic feature. This is the “degree” of each locus in a network where each TF bound locus or ChAR is a node, and an edge between nodes exists if linked by Batf HiChIP loops. **(C)** Percent of regions with row feature (loci bound by TF or ChAR gained and lost in Batf cKO and Irf4 cKO) with a long-range HiChIP loop connected to column feature. Loop length threshold = 1,000,000 bp. Enrichment > 0.4 are highlighted with black edges. **(D)** Percent of regions that interact with DE genes linked by Batf long-range loops. A loop was considered to contact a DE gene if one of its loop anchors spanned the promoter of the gene. **(E)** Representative long-range contacts/loops in WT CD8+ T cells around *Ctla4* and *Icos* loci identified by HiChIP for Batf (red arc) and Ctcf (blue-green arc). Coordinates of ChIP-seq peaks identified by MACS2 for Batf are displayed as black bars.

Taking a closer look, we found that regions that were bound by certain TFs tended to form larger than expected numbers of long-range interactions (Figure **<u>3B</u>**; Wilcoxon test adjusted p-value < 0.05 on each test comparing regions with a given feature to those without). This tendency was especially pronounced in regions bound by Runx3 or T-bet, and in “hot” regions that are occupied by many TFs (resonating with the notion of super enhancers (Huang et al., 2018)). Finally, through pairwise analysis of our data, we found that many TF binding sites came into contact with one another or with Batf-dependent ChARs through Batf-mediated loops (Figure **<u>3C</u>**). These trends may be reflective of the effects of Batf on transcription, as we observed many cases by which key regions (TF binding sites, or Batf-dependent ChARs) come into contact with the promoters of genes with a Batf-dependent expression level (Figure **<u>3D</u>**). As an example, we observed a dense set of Batf-anchored loops around the *Ctla4* and *Icos* loci, which were both downregulated in Batf cKO (Figure **<u>3E</u>**). These loops brought in close proximity a group of regulatory regions that required Batf for accessibility and were bound by most of the TFs we examined (Figure **<u>S3A</u>**).

Taken together, these results associate many of the observed effects of Batf on the transcriptome and epigenome with higher-order organization of the chromatin. Specifically, they suggest that Batf-dependent chromatin loops may serve to bring in proximity regions that are bound by key TFs, and that Batf may be critical for both the accessibility of these regions and for the long range contacts they make. Among the TFs we studied, the extent of overlap was most pronounced for Runx3 and T-bet, with many binding sites coinciding with Batf mediated loops (Figure **<u>3A</u>**), forming larger numbers of loops per-site (Figure **<u>3B</u>**). Furthermore, Runx3 and T-bet exhibited the largest proportion of Batf-mediated interactions with regions bound by other TFs and with regions with Batf-dependent accessibility (Figure **<u>3C</u>**). These results thus suggest a strong link between Batf-mediated organization of the chromatin and the activity of Runx3 and T-bet.

### Overexpression of Batf and other key transcription factors recapitulates aspects of CD8+ T cell chromatin landscape and gene expression in fibroblasts

Having observed that Batf and Irf4 affect critical chromatin features that are required for CD8+ T cell differentiation and function, and that T-bet and Runx3 preferentially bind regions that are regulated by Batf and Irf4, we sought to determine whether these four transcription factors are sufficient for recapitulating CD8+ T cell chromatin and transcriptional properties in a different cell type. Thus, we set up an ectopic gene expression system in NIH/3T3 fibroblasts, which allowed us to express all possible combinations of these four TFs, and measure the resultant effects on chromatin accesiblity and the transcriptome.

We first used western blot and RNA-seq to confirm that the endogenous expression of Batf, Irf4, Runx3 and T-bet in NIH/3T3 fibroblasts was undetectable (Figure **<u>S4A</u>** and **<u>4A</u>**). We then transduced the fibroblasts with a doxycycline-inducible lentiviral vector consisting of any combination of Batf, Irf4, Runx3 and T-bet (16 conditions altogether). Successfully transduced cells were selected with antibiotics or fluorescence (Figure **S4B)**. Ectopic expression of the respective TFs in each sample was then induced simultaneously with doxycycline for 72 hours, as confirmed by western blots and RNA-seq (Figure **<u>S4A</u>** and Figure **<u>4A</u>**). Transduced fibroblasts were subjected to profiling with RNA-seq (<u>Table **6**</u>) and ATAC-seq (<u>Table **7**</u>), providing an estimate for the changes in the transcriptome and chromatin landscape for each combination of TFs. As a negative control we also assayed the same set of transcription factor combinations with no doxycycline induction (16 control conditions altogether).

**Figure 4.**
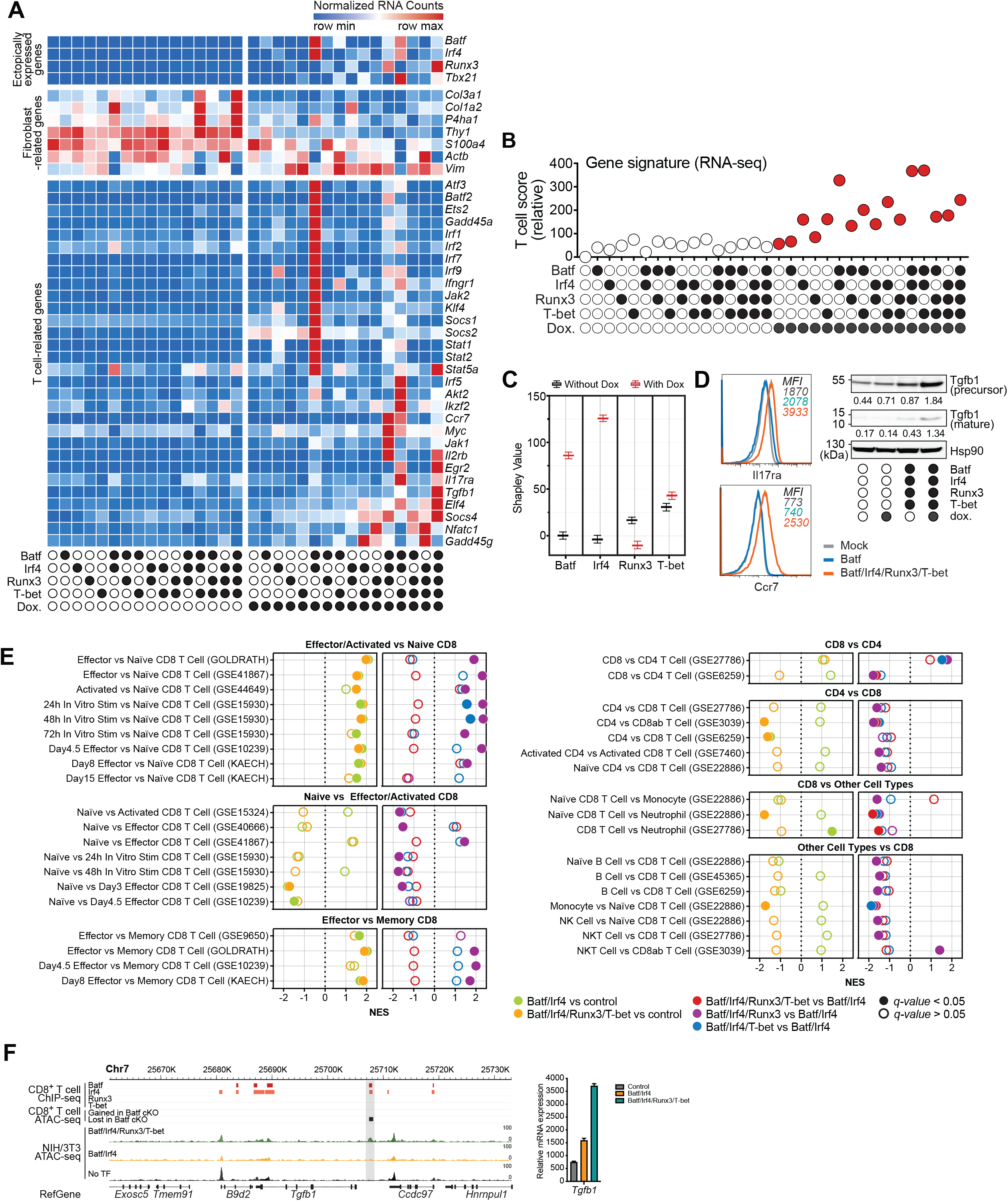
Overexpression of Batf and other key transcription factors recapitulates aspects of CD8 T cell chromatin landscape and gene expression. **(A)** Row normalized gene expression of selected T cell and fibroblast genes co-expressing four TFs combinations with or without doxycycline induction. **(B)** Relative T cell score of normalized gene expression for all experimental conditions in fibroblast. See **Methods**. **(C)** Box and whisker plot of the Shapley value for each TF, based on the relative T cell score. **(D)** Validation of selected surface markers and protein expression by flow cytometry and western blot quantified with densitometry for doxycycline-treated fibroblasts expressing no transcription factors, Batf only or Batf/Irf4/Runx3/T-bet. **(E)** Normalized Enrichment Score (NES) for pathways found in GSEA based on differential expression results in fibroblasts. The conditions tested are indicated by color and include fibroblasts expressing Batf/Irf4, Batf/Irf4/Runx3/T-bet, or no TFs controls. Gene sets and immunologic signatures related to CD8+ T cells are shown. **(F)** Representative tracks showing the chromatin accessibility changes in fibroblasts around *Tgfb1* loci, when expressing Batf/Irf4/Runx3/T-bet, Batf/Irf4 and no TFs. Batf, Irf4, Runx3, T-bet ChIP-seq peaks in CD8+ T cells and DA ChARs from Batf cKO are also shown as references for TF binding and chromatin accessibility. Shaded areas indicated putative regulatory regions for *Tgfb1* in fibroblasts. Bar plot of the relative mRNA expression of *Tgfb1* from control, Batf/Irf4 and Batf/Irf4/Runx3/T-bet expressing fibroblasts is shown on the right.

Focusing on the gene expression data, we found that different combinations of the four TFs led to increased expression of several genes involved in the regulation of T cell homeostasis, activation, proliferation, differentiation and function (Figure **<u>4A</u>**), such as IRF family members (*Irf1*, *Irf2*, *Irf5*, *Irf7* and *Irf9* (Hammami et al., 2015; Hida et al., 2000; Martinet et al., 2015; Simon et al., 1997; Zhou et al., 2012)), JAK-STAT signaling pathway members (*Jak1*, *Jak2*, *Stat1*, *Stat2* and *Stat5a* (Seif et al., 2017)), SOCS family members (*Socs1*, *Socs2* and *Socs4 (Palmer and Restifo, 2009)*), cytokine signaling (*Tgfb1*, *Il2rb*, *Ifngr1* and *Il17ra* (Curtsinger et al., 2012; Oh and Li, 2013; Ross and Cantrell, 2018; Tosello Boari et al., 2018)) and other factors (*Atf3*, *Batf2*, *Ets2*, *Gadd45a*, *Klf4*, etc. (Filén et al., 2010; Kayama et al., 2019; Salvador et al., 2005; Yamada et al., 2009; Zaldumbide et al., 2002)). To more systematically evaluate these transcriptional effects, we created a gene signature consisting of genes that are the most differentially expressed (up or down) between untreated fibroblasts and CD8+ T cells (using DESeq2; see **Methods**). We then used a method akin to Vision (DeTomaso et al., 2019) to calculate a signature score for each condition, estimating its position in the spectrum between CD8+ and fibroblast transcriptional states (Figure **<u>4B</u>**). We observed that the ectopic expression of Batf and Irf4 is generally associated with a high T cell score (closer to T cell state than others). Conversely, ectopic expression of Irf4 only or Batf only resulted in less pronounced effects, supporting the model of synergism between these two TFs.

To assess each TF’s individual contribution to changes in our T cell score, we used the Shapley Value - a game theoretic measure that summarizes the contribution of individuals in cooperatie scenarios (Keinan et al., 2004). Batf and Irf4 scored the highest, indicating that these two TFs played the primary role in reshaping the transcriptome of fibroblasts toward a T cell-like profile (Figure **<u>4C</u>**). The score for T-bet was substantially lower, and Runx3 had a slightly negative score. This indicates that these two TFs had smaller impacts from the genome-wide perspective, but not excluding a more nuanced contribution in regulating specific genes.

Indeed, while over expression of Batf and Irf4 resulted in marked increase in the expression of critical T cell factors (Figure **<u>4A</u>**), we found parts of the T cell program that were most strongly upregulated when Runx3 and/or T-bet were ectopically expressed alongside Batf and Irf4, including chemokine receptors such as *Ccr7*, cytokines and cytokine receptors such as *Il2rb, Il17ra* and *Tgfb1*, and T cell transcription factors *Stat5a*, *Egr2* and *Elf4* ( Figure **<u>4A</u>**, <u>Table **6**</u>). To confirm this, we selected genes that play important roles in CD8+ T cell homeostasis, survival and function (Acharya et al., 2017; Gaffen, 2009; Jung et al., 2016; Tinoco et al., 2009; Tosello Boari et al., 2018) and examined their expression with flow cytometry (*Il17ra* and *Ccr7*, Figure **<u>4D</u>** left) or western blot (*Tgfb1*, Figure **<u>4D</u>** right). Consistent with our RNA-seq analysis, we observed that these genes become induced in the treated fibroblasts and that their expression is most pronounced when all four TFs are present. To gain further insight into these transcriptional changes, we applied GSEA using as reference the literature-curated MSigDB C7 collection of transcriptional dissimilarities between subsets of immune cells (Godec et al., 2016) (Figure **<u>4E</u>**, <u>Table **8**</u>). Overall, we found that ectopic expression of Batf and Irf4 resulted in transcriptional differences akin to those between effector CD8+ T cells and other CD8+ T cell states (e.g. naive and memory; Figure **<u>4E</u>**, left). In agreement with most of the immune signatures we listed, the addition of Runx3, T-bet or both further strengthen the differences (compared to fibroblasts treated with only Batf and Irf4) between effector CD8+ versus other CD8+ populations. Runx3, T-bet or both TFs were also observed to differentiate the CD8+ identity when comparing CD8+ signatures with other cell types (e.g. CD4+, B cell, monocyte, NK and NKT cells; Figure **<u>4E</u>**, right). Taken together, this analysis indicates that over expression of the four TFs resulted in a shift toward the effector identity within the CD8+ T cell populations.

Given the role of Batf in modulating the epigenome of CD8+ T cells, we reasoned that the transcriptional changes we observed are accompanied by epigenetic reprogramming of the treated fibroblasts. Indeed, signature analysis of global changes in chromatin accessibility with ATAC-seq (using as signature regions that are differentially accessible between fibroblasts and T cells, in a manner akin to Figure **<u>4B</u>**) points to a similar phenomenon, with gradual modulation of the chromatin accessibility landscape in fibroblasts toward T cell like profile (Figure **<u>S4C</u>**). As an example, we considered loci around *Tgfb1* (Oh and Li, 2013), which is not normally expressed by fibroblasts but became induced in our ectopic system (Figure **<u>4F</u>**). We observed a specific locus downstream of *Tgfb1* that became accessible only when the four TFs were expressed. This locus is also associated with loss of accessibility in Batf cKO and with direct binding by Batf and Irf4 in T cells (Table **2**, Figure **<u>S5A</u>**) and may therefore act as an enhancer of the nearby *Tgfb1* locus, whose activity in T cells depends on all four TFs.

Taken together, our results suggest a major role for Batf and Irf4 in shaping the transcriptome and epigenome of CD8+ T cells, due to their effects on a subset of key genes and loci that are otherwise inactive in fibroblasts. The results also suggest that Runx3 and T-bet may be important for fine-tuning or enhancing these global effects, toward a more specific effector CD8+ T cell like phenotype.

### Establishment of long range chromatin loops characteristic of T cells requires overexpression of Runx3 and T-bet in addition to Batf and Irf4

Having found that Batf bound loci engage in long-range genomic contacts in T cells, we sought to determine if its ectopic expression (alone or with its associated factors) can reconstitute similar chromatin structures in fibroblasts. To this end, we performed Batf HiChIP experiments in fibroblasts expressing only Batf or a combination of all four transcription factors.

Examples of Batf-anchored chromatin loops in these manipulated fibroblasts around the *Nfatc1* and *Tgfb1* loci are shown in figure **<u>5A</u>** and **<u>S5A</u>**. Ectopic expression of Batf alone generated mostly short range loops near these loci, which only partially recapitulated the Batf loops in WT effector CD8+ T cells. However, when Batf/Irf4/Runx3/T-bet were co-expressed, the resulting Batf-anchored loops recapitulated many of the loops found in T cells, especially long-range contacts. Noticeably, the 5’ proximal promoter region of *Nfatc1* has a Batf-dependent accessibility and is co-bound by Batf and Irf4 in T cells. It is also linked by Batf- mediated loops to the 3’ end of that gene, which (in T cells) includes binding sites for Batf, Irf4, Runx3 and T-bet. Finally, we note that Batf alone did not alter the expression of *Nfatc1* in fibroblasts, which required the addition of other factors (Figure **<u>4A</u>**). These findings therefore point to a composite epigenetic structure, (TF binding, chromatin accessibility and chromatin loops) that may contribute to the expression of a nearby gene, while depending (directly or indirectly) on the joint activity of several TFs, in addition to Batf.

**Figure 5.**
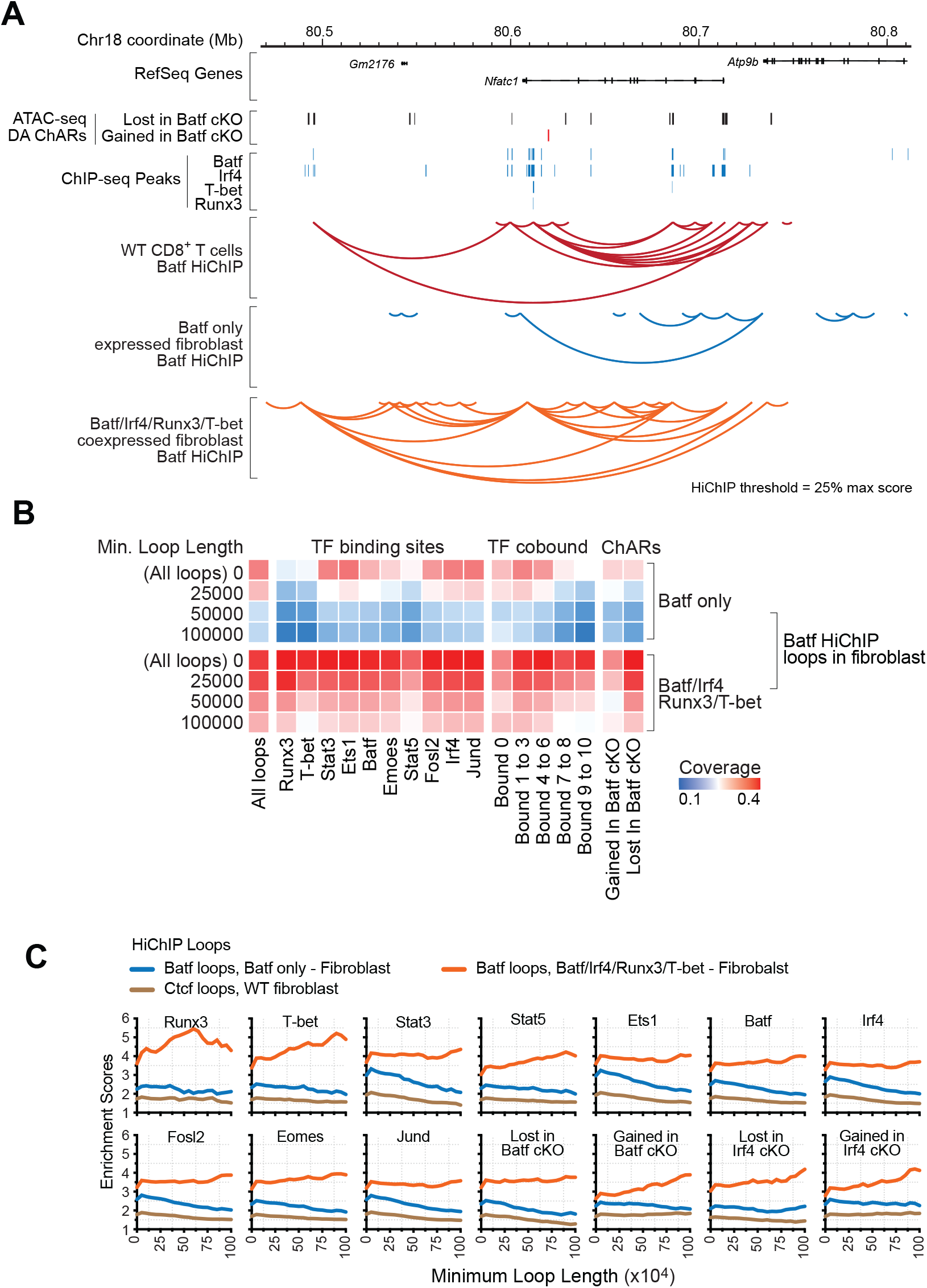
Overexpression of Batf, Irf4, Runx3 and T-bet are able to reconstruct chromatin architectures of NIH/3T3 fibroblasts. **(A)** Representative long-range genomic contacts/loops around *Nfatc1* locus identified by HiChIP for Batf in WT CD8+ T cells (red arcs), Batf in fibroblasts expressing Batf (blue arcs), and Batf in fibroblasts co-expressing Batf/Irf4/Runx3/T-bet (orange arcs). DA ChARs identified in WT vs Batf cKO CD8+ T cells are shown as red and blue bars (lost and gained in Batf cKO, respectively). ChIP-seq peaks identified for Batf, Irf4, Runx3 and T-bet in WT CD8+ T cells are displayed in black bars. **(B)** Coverage (percent of Batf HiChIP loops where both loops anchors are overlapped by both anchors of at least one loop in fibroblasts expressing Batf, or fibroblasts expressing Batf/Irf4/Runx3/T-bet) for different lengths of loops overlapped with defined features, including all loops, TF binding sites and DA ChARs in Batf cKO. **(C)** Fold enrichment of area spanned by HiChIP loop anchors by HiChIP for Batf in fibroblasts expressing Batf (blue), Batf in fibroblasts co-expressing Batf/Irf4/Runx3/T-bet (orange) and Ctcf (brown) in WT fibroblasts in areas spanned by key CD8+ TFs and ChARs gained/lost in Batf cKO and Irf4 cKO CD8+ T cells. Fold enrichment is defined as “percent of genome spanned by feature / percent of HiChIP loop anchor area spanned by feature”.

A more global comparison of Batf-dependent genomic contacts in T cells vs. manipulated fibroblasts indeed reveals a higher level of concordance in cells co-expressing Batf/Irf4/Runx3/T-bet vs. Batf alone, and with differences most pronounced in the case of long-range loops (Figure **<u>5B</u>** and **<u>S5B</u>**). Indeed, overall we observed that approximately 40% of the Batf-dependent loops in T cells were recapitulated in fibroblasts when overexpressing all four TFs versus 20% when over expressing Batf alone. The increase in concordance was observed more specifically next to regions that were functionally important in CD8+ T cells, namely Batf-dependent ChARs (especially ones that lose accessibility in Batf cKO), Irf4-dependent ChARs, and regions bound by key TFs (Figure **<u>5B</u>** and **<u>5C</u>**, <u>Table **10**</u>). This trend was especially strong in regions bound by Runx3, T-bet, Stat3, and Ets1 in T cells, or regions occupied by many TFs (Figure **5B, 5C** and **<u>S5B</u>**).

Taken together, these results demonstrate that overexpressing the four TFs, Batf, Irf4, Runx3 and T-bet in fibroblasts, but not Batf alone, was sufficient to reconstitute a chromatin architecture that was closer to that observed in CD8+ T cells. It therefore supports the notion that Batf exerts its regulatory effects, at least in part, by modulating the long range architecture of the chromatin, and that these changes required the presence of other cofactors, including Irf4, Runx3 and T-bet.

## DISCUSSION

The differentiation of naive CD8+ T cells into an effector state is a critical step in adaptive immunity. Although many of the TFs that govern T cell differentiation or response to stimulation have been characterized, the regulation of the earliest epigenetic and transcriptional events that initiate the transition of cytotoxic T cells into an effector state remains partially understood. Previous work by us and others (Godec et al., 2015; Grusdat et al., 2014; Kurachi et al., 2014; Kuroda et al., 2011) identified the transcription factor Batf to be essential for establishing commitment into the effector state. Here, we further explored how Batf, along with other key TFs, may achieve its influence on this process.

Our study provides a detailed account of the transcriptome and epigenome of differentiating CD8+ T cells in mice, and their dependence on the proper functioning of Batf. Our data set includes genome-wide characterization of the transcriptome of CD8+ T cells (RNA-seq) in an *in vivo* infection model and in a time course of naive CD8+ T stimulated *in vitro*. It also includes a map of the accessible chromatin landscape (ATAC-seq), binding of ten key TFs (ChIP-seq) and mapping of long-range chromatin loops that are anchored by Batf (Hi-ChIP). To probe the causal effects that Batf may have on the differentiation and function of CD8+ T cells, we repeated our primary measurements using samples from Batf KO animals. Joint analysis of these data resulted in a map of the CD8+ T cell epigenome, consisting of 96,490 genomic regions along with the TFs that bind them, their accessibility, their contacts with other regions, the expression from nearby genes, and the dependence of all these modalities on the presence of Batf. Through this joint analysis, we observed that the absence of Batf had a substantial effect on both the transcriptome and the structure of the chromatin. The transcriptional effects, which were observed in thousands of genes, largely indicated that Batf contributes to a shift into an effector state, whereby KO cells tend to retain a naive-like gene expression program with markers such as *Sell* and *Ccr7*. The respective epigenetic changes were often observed in proximity to the affected genes and had a similar level of abundance, with thousands of affected loci. Intriguingly, we observed these effects on several levels: the increase or decrease in accessibility often coincided with changes to binding of key TFs, which in turn were observed close to Batf-anchored chromatin loops.

The availability of genome wide estimates of both chromatin accessibility and protein-DNA interactions can lend insight into the manner by which Batf affects different TFs. In most cases, we observed a positive correlation between changes in binding and changes in accessibility (comparing KO to WT cells). While it may suggest a simple global mechanism by which Batf-dependent accessibility is required for binding, we cannot rule out other alternatives. Indeed, it is interesting to note that for some TFs (most prominently T-bet, Stat5 and Jund), changes to binding were restricted to decrease, which became more severe as accessibility decreased. Conversely, in other TFs (e.g., Runx3, Irf4) we also see an increase in binding at loci that became more accessible in the absence of Batf. While Batf- dependent changes to the mRNA expression of these TFs might in principle serve to explain such differences, we have not observed such trends in our data (Figure **<u>S6D</u>**). It is therefore conceivable that our observations may result from additional (direct or indirect) consequences of Batf function, such as changing the expression of other TFs outside our panel, modifications of TF proteins or nearby histone tails, and local alterations in binding affinity through protein-protein contacts. The latter option is further illustrated in the case of Fosl2 and Stat3. Unlike the other transcription factors we assayed, Fosl2 and Stat3 exhibited marked increase in binding in Batf KO, but this increase had only weak association with accessibility. While there may again be several alternative explanations, it is interesting to note that in both cases the increase in binding may result from reduced activity of a binding competitor (Batf and Stat5, respectively).

The observation that Stat5 lost its binding while Stat3 gained binding in the absence of Batf further suggests a general regulatory mechanism that controls the activity of Stat family proteins. Stat3 and Stat5 are known to be activated by the IL-2 signaling pathway, however while they have been demonstrated to be non-redundant and function differently in T cells (Laurence et al., 2007; Moriggl et al., 1999), the mechanisms that balance their usage are not well characterized. Our data therefore suggests an additional function of Batf that may serve to regulate the activity of these two proteins. Within our system, this balancing raises an hypothesis as to what may contribute to the enhanced T cell death and subsequent lack of functional effector cells in infected naive mice transferred with Batf KO cells (Kurachi et al., 2014). Although Batf-deficient cell counts diminished on day 4 after LCMV challenge, the KO cells can be easily cultured and expanded *in vitro* in the presence of recombinant IL-2. Given that the expression of IL-2 receptors (*Il2ra, Il2rb and Il2rg*) were intact in Batf cKO (Figure **<u>S6A</u>**), the defects of Stat5 binding and increased Stat3 binding in Batf cKO (Figure **<u>S6B</u>**) may contribute to the IL-2 unresponsiveness and cell death observed in Batf KO since Stat5 is the immediate downstream signal transducer of IL-2 signaling pathway (Lin et al., 1995). Our data points to additional (possibly downstream) potential mediators of the observed cell death in Batf KO in vivo, which may rely on some, but not all, of our measurement modalities. For instance, the apoptotic factor Bcl2l11 (*Bim*) (Bouillet et al., 2002) has elevated expression in the absence of Batf. We further find that its promoter comes in Batf-mediated interaction with a distal site (> 300 kb), and that both promoters and distal sites are bound by Batf, Irf4, Runx3, and T-bet (Figure **<u>S6C</u>**). However, the accessibility of these sites and the majority of their binding events are not dependent on Batf, raising the option of involvement by additional factors that are not included in our measurements.

In agreement with previous observations of similar phenotypes in mice deficient for either Batf or Irf4 (Grusdat et al., 2014), we found that perturbing these two proteins resulted in similar trends in terms of the transcriptome and epigenome. These effects, however, were not completely overlapping. For instance, while the absence of Batf had a significant effect on the expression of interferon signaling genes, this effect was not as strongly observed with Irf4. More globally, the number of regions differentially accessible in Batf cKO was almost twice as much as in Irf4 cKO, suggesting that it plays a broader or more critical role in reshaping the chromatin post stimulation. Either Batf or Irf4 alone, however, were insufficient for inducing broad changes in the epigenome of fibroblasts, while their joint effect was sufficient to engage some aspects of a CD8+ core program, thus adding to a growing body of evidence on cooperativity between these two molecules (Grusdat et al., 2014; Man et al., 2017; Yao et al., 2013).

In addition to Irf4, cumulative evidence from all of our genomic assays pointed to two additional TFs, Runx3 and T-bet, as key regulators which may work in concert with Batf and Irf4. Specifically, we found that Runx3 and T-bet tend to bind in regions whose accessibility depends on Batf and that the strength of binding changed proportionally to Batf-dependent changes in accessibility. They also had the strongest tendency to bind next to other transcription factors (particularly Batf) through direct genomic proximity or through long-range Batf-mediated loops. Finally, both Runx3 and T-bet tended to be connected, by long-range chromatin loops, to the promoters of genes with a Batf-dependent expression. Indeed, ectopic expression in fibroblasts provided evidence that these factors contribute to strengthening the induction of a T-cell like transcriptional program, when co-expressed with Batf and Irf4.

These results accord with reports that T-box proteins and Runx3 contribute cooperatively to the effector program of CD8+ T cells (Cruz-Guilloty et al., 2009). They further agree with more recent publications (van der Veeken et al., 2019; Wang et al., 2018), which showed that T-box factors and RUNX family proteins are required to drive and maintain the changes in the chromatin landscape during effector differentiation and memory generation of CD8+ T cells. Adding to that, our over-expression analysis suggests that T-box and Runx3 may act synergistically with Batf and Irf4 to induce a T-cell like program, in terms of gene expression, chromatin accessibility and long range chromatin loops. The mechanism behind these effects is yet to be determined though. While it has been observed that Runx3 and T-bet can physically interact with each other(Djuretic et al., 2007), currently there is no evidence showing physical associations with Batf or Irf4. One study showed the interaction between AP-1 (Fos and Jun) and Runx family proteins(D’Alonzo et al., 2002) which raised the possibility of Batf interacting with AP-1, Runx3 and T-bet in the same complex, since Batf interacts with AP-1 constantly. Interestingly, however, crystal structure study showed that T-bet was capable of binding to two independent strands of DNA and may function as a chromatin linker(Liu et al., 2016). Thus, Batf may connect the co-regulated regions through the binding of T-bet to form a regulatory complex.

While our focus was on the cooperation between Irf4, Runx3, T-bet and Batf in effector T cells, our data may be further explored for insight on the activity of other TFs, and possibly on the regulation of other CD8+ T cell states. Indeed, Batf has been shown to also be expressed by naive and exhausted T cells, and may interact with different proteins in different cell contents (Chang et al., 2014; Man et al., 2017). For instance, while the expression of T-bet was shown to characterize naive and effector T cells, the activity of Eomes, a T-box TF with high homology to T-bet which we profiled here as well, has been more prominently associated with memory and exhausted T cells (Buggert et al., 2014; Sen et al., 2016). The extensive overlap we observed between Batf and Eomes binding sites, their proximity to genes and their dependence on Batf may therefore provide further hypotheses for regulation in these subsets.

## SUPPLEMENTARY FIGURE LEGENDS

**Supplementary Figure 1. (A)** Schematic of Cre-lox conditional knockout system and lymphocytic choriomeningitis virus (LCMV) acute infection mouse model. **(B)** Venn diagram showing the overlap between RNA-seq DE genes (WT vs Batf cKO, both up-regulated and down-regulated) and the genes associated with DA ChARs in ATAC-seq (WT vs Batf cKO, both up-regulated and down-regulated, consider only the nearest gene defined by GREAT). Hypergeometric p-value for the overlap is shown. **(C)** Scatter plot showing the relationship between chromatin accessibility and transcriptome changes in Batf cKO vs WT CD8+ T cells. Only genes/loci which are significant at adjusted p-value <0.05 in either test displayed. **(D)** Representative Tn5 pileup from ATAC-seq at differential loci (*Sell* and *Maf*) identified by DESeq2. Relative mRNA expression levels of *Sell* and *Maf* from RNA-seq in Batf cKO and WT CD8+ T cells are shown as barplots. **(E)** Volcano plot of log2 fold change (Irf4 cKO/WT) in gene expression. **(F)** Volcano plot of log2 fold change (Irf4 cKO/WT) in chromatin accessibility (Tn5 cuts) at loci identified as accessible (MACS2 peak) in either Irf4 cKO or WT conditions. Name of the nearest gene mapped by GREAT is displayed for a subset of loci. **(G)** PCA plot of cuts in ATAC-seq peaks of naive, Batf cKO, Irf4 cKO and control CD8+ T cells in LCMV infection model. **(H)** DE gene counts of WT vs Batf cKO or Irf4 cKO. **(I)** DA ChAR counts of WT vs Batf cKO or Irf4 cKO. **(J)** GSEA of DE genes from Batf cKO and Irf4 cKO CD8+ T cells. Selected gene signatures and clusters of gene expression from in vitro culture are shown. Only results with adjusted p-value < 0.05 are displayed. **(K)** Fold enrichment for selected immunologic signatures of WT vs Irf4 cKO DA ChARs GREAT pathways. WT DA ChAR-associated gene sets are shown in red and Irf4 cKO DA ChAR-associated gene sets are shown in blue. **(L)** Top 50 enriched transcription factor motifs in DA ChARs identified in Batf cKO CD8+ T cells with JASPAR database (Khan et al., 2017), by fold-enrichment in peaks differentially open in Batf cKO or WT conditions relative to union of peaks across the two conditions. Red circles represent DA ChARs enriched in WT (lost in cKO) and blue circles represent DA ChARs enriched in Batf cKO (lost in WT). Filled circles are datasets with an adjusted p-value < 0.05 whereas open circles are datasets with > 0.05.

**Supplementary Figure 2. (A)** Representative pileup of ChIP-seq fragments from Batf KO and WT CD8+ T cells (normalized by control sample in MACS2) for key transcription factors (transcription factors) around selected genes. **(B)** Fold enrichment of transcription factor binding over the DA ChARs lost in Irf4 cKO samples. Off-diagonal entries in the heatmap show the enrichment of regions co-bound by the two respective transcription factors. Diagonal entries in the heatmap and the barplot to the right show the enrichment for individual transcription factors. See methods section for details of fold enrichment calculation.

**Supplementary Figure 3. (A)** Long-range contacts/loops in WT CD8+ T cells around *Ctla4* and *Icos* loci identified by HiChIP for Batf (red arc) and Ctcf (blue-green arc). Coordinates of ChIP-seq peaks identified by MACS2 for indicated transcription factors from WT CD8+ T cells are displayed as blue bars. The DA ChARs identified in ATAC-seq datasets for regions that are lost and gained in Batf cKO cells are shown as black and red bars, respectively.

**Supplementary Figure 4. (A)** Western blot validating protein expression of the four ectopically expressed CD8+ TFs across all experimental conditions.. Heat shock protein 90 (Hsp90) was used as internal control. **(B)** Top: constructs of the doxycycline inducible TF expression system. HPH: Hygromycin B phosphotransferase, PAC: Puromycin N-acetyltransferase, BSD: Blasticidin-S deaminase. Bottom: schematic of lentiviral transduction and doxycycline inducible system for expressing four key CD8+ T cell TFs in fibroblasts. Chart was created with BioRender.com **(C)** Percentage of T cell score of normalized DNA accessibility for all experimental conditions. T cell accessibility signatures obtained from a DESeq2 comparison of ATAC-seq of effector and naive CD8+ T cells versus fibroblasts. Signature is defined by giving a weight of +1 to the top 1000 peaks differentially upregulated in T cells and weight of −1 to the top 1000 peaks differentially upregulated in fibroblasts. These weights were then applied to the vector of normalized counts for each condition.

**Supplementary Figure 5. (A)** Representative long-range genomic contacts/loops around *Tgfb1* locus identified by HiChIP for Batf in WT CD8+ T cells (red arcs), Batf in fibroblasts expressing Batf (blue arcs), and Batf in fibroblasts co-expressing Batf/Irf4/Runx3/T-bet (orange arcs). DA ChARs lost in WT vs Batf cKO CD8+ T cells are shown as blue bars. ChIP-seq peaks identified for Batf, Irf4, Runx3 and T-bet in WT CD8+ T cells are displayed in black bars. **(B)** Jaccard statistics assessing recovery of T cell loops in fibroblasts expressing Batf or Batf/Irf4/Runx3/T-bet. Statistics are stratified by epigenetic features found on either anchor of the loop. Minimum loop length was set to 0.

**Supplementary Figure 6 (A)** Relative mRNA levels of *Il2ra*, *Il2rb* and *Il2rg* of WT, Batf cKO and Irf4 cKO CD8+ T cells sorted from day 3.5 LCMV challenged mice. **(B)** Representative IGV pileup of Stat3 and Stat5 ChIP-seq fragments from CD8+ T cells culture *in vitro* around *Bim* (*Bcl2l11*) locus, both WT and Batf KO. **(C)** Representative plot (adapted from WashU Epigenome Browser) to show the ChIP-seq peaks (Batf, Irf4, Runx3 and T-bet from WT CD8+ T cells), DA ChARs identified with MACS2 (WT vs Batf cKO), merged regulatory regions and HiChIP (Batf, WT CD8+ T cell) loops around *Bim (Bcl2l11)* locus. **(D)** Normalized RNA counts of transcription factors from WT and Batf cKO CD8+ T cells sorted from LCMV mouse model.

## SUPPLEMENTARY TABLE LEGENDS

**Table 1 - T cell RNA-seq**

This table lists our RNA-seq experiments in T cells (in vitro and in vivo), and reports the normalized counts for each experiment as well as the differential expression results.

**Table 2 - T cell ATAC-seq and ChIP-seq**

Each row in this table corresponds to a locus in the genome. Each column indicates an in vitro ChIP-Seq experiment or in vivo ATAC-Seq (listed in the first row), and indicates whether a peak was found with MACS2 and whether it was differentially accessible (ATAC-Seq) or differentially bound (ChIP-Seq) between WT and KO conditions.

**Table 3 - T cell GSEA**

Gene set enrichment analysis (GSEA) results for differentially expressed (DE) genes, comparing knockouts (Irf4 or Batf) to WT CD8+ T cells. The underlying DE genes can be found in Table 1.

**Table 4 - T cell chromatin enrichment analysis**

This table reports whether the T cell chromatin features in Table 2 were found to be associated with transcription factor binding motifs, disease-associated SNPs, and other genomic annotations, and includes analysis from GREAT. See section “**Locus- level enrichment analysis”** in methods for details.

**Table 5 - T cell HiChIP**

This table reports the summary statistics for the T Cell HiChIP experiments.

**Table 6 - Fibroblast RNA-Seq**

This table reports differential expression results and normalized counts for the in vitro fibroblast experiments.

**Table 7 - Fibroblast ATAC-seq**

Each row in this table corresponds to a locus in the genome. Each column indicates an in vitro ATAC-Seq (listed in the first row) experiment, and indicates whether a peak was found with MACS2 and whether it was differentially accessible under different experimental conditions.

**Table 8 - Fibroblast GSEA**

Gene set enrichment analysis (GSEA) results for fibroblast experiments. The underlying DE results can be found in Table 6.

**Table 9 - Fibroblast chromatin enrichment analysis**

This table reports whether the fibroblast ATAC-seq peaks from different experimental conditions in Table 7 were found to be associated with transcription factor binding motifs, disease-associated SNPs, and other genomic annotations, and includes analysis from GREAT. See section “**Locus-level enrichment analysis”** in methods for details.

**Table 10 - Fibroblast HiChIP**

This table reports the summary statistics for the fibroblast HiChIP experiments.

## SUPPLEMENTARY RESOURCES

Our epigenetic datasets (ATAC-Seq peaks, ChIP-Seq peaks, and HiChIP loops) are available for browsing on the WashU Epigenome Browser (Li et al., 2019) at https://epigenomegateway.wustl.edu/browser/?genome=mm10&sessionFile=https://yosef-lab-public-resources.s3-us-west-1.amazonaws.com/BATF/WashU_Browser/Datahub/BATF_Session_All_Tracks.json.

## MATERIALS AND METHODS

### Mice and LCMV infection model

Batf KO mice in B6 background were purchased from the Jackson Laboratory (Schraml et al., 2009) and crossed with P14 TCR transgenic mice to generate P14 Batf KO mice (Kurachi et al., 2014). Five to 10 weeks of age female mice were used in this study. All animal work was performed in accordance with the Institute Animal Care and Use Guidelines of the University of Pennsylvania.

For the generation and adoptive transfer of Batf cKO P14 CD8+ T cells, CD45.2^+^ Batf-floxed and Irf4-floxed ERT2-Cre P14 donor female mice was treated with 2 mg 4-hydroxytamoxifen (4-OHT) per day per mouse intraperitoneally for 5 consecutive days. After the Cre-mediated recombination of the loxP flanked sequence, Batf was removed along with GFP. For Irf4 conditional KO constructs, Irf4 followed by a stop signal was excised by the induced Cre activity, thus inducing expression of YFP. Five days after the final dosing, mice spleens were then harvested and Batf cKO and Irf4 cKO CD8+ P14 T cells were sorted with cell sorter (BD, FACSAria) according to the GFP and YFP fluorescence expression, respectively. The sorted Batf cKO and Irf4 cKO CD8+ T cells were mixed with WT P14 CD8+ T cells (2 ×10^4^) and adoptively transferred into WT mice expressing CD45.1 congenic marker.

LCMV Armstrong strain was produced and viral titers were quantified as described previously (Kao et al., 2011). Mice were challenged by intraperitoneal injection (i.p.) of LCMV with 2 × 10^5^ plaque forming units (PFU) one day after the adoptive transfer.

### In vitro CD8+ T cell culture

Spleens from WT and Batf KO P14 mice were harvested, homogenized and passed through 70 um cell strainers. Splenocytes were spun down at 500 g for 10 min at 4 °C in 50 ml centrifuge tubes and the red blood cells were lysed with 1 ml ACK lysis buffer per spleen for 1 min at room temperature. Cells were washed once with MACS buffer (1X PBS, 0.5% BSA, 2 mM EDTA) and cell numbers were counted. Mouse naive CD8a+ T cells isolation kit (Miltenyi Biotec, 130-096-543) was used for CD8+ T cells isolation according to manufacturer’s instructions. The isolated CD8+ T cells were plated on anti-CD3 (BD Pharmingen, 553057, 2 ug/ml in PBS, coating at 4°C overnight) pre-coated 24-well plates with the concentration of 1 × 10^6^ cells/well in 1 ml complete RPMI medium (RPMI with 1X NEAA, 10 mM HEPES, 1 mM sodium pyruvate, 10% FBS, 55 mM 2-Mercaptoethanol, 50 U/ml Penicillin and 50 ug/ml Streptomycin) supplemented with 100 U/ml hIL-2 and 2 ug/ml anti-CD28 (BD Pharmingen, 553294). Two days after the primary stimulation, 1 ml fresh complete RPMI was added to the same well with 100 U/ml hIL-2. The cultured CD8+ T cells were split and expanded into larger culture wells or flasks every day from day 3 with fresh RPMI medium with 100 U/ml hIL-2 until day 6. On culture day 6, cells were re-stimulated with 50 ng/ml Phorbol 12-myristate 13-acetate (PMA, Sigma P1585) and 1 uM ionomycin (Sigma, I0634) for additional 3 hours and harvested for further experiments.

### RNA sequencing

Total RNA was extracted with QIAGEN RNeasy Plus Mini Kit (QIAGEN, 74134). mRNA was isolated with NEBNext Poly(A) mRNA Magnetic Isolation Module (NEB, E7490) and the RNA-seq libraries were generated with NEBNext Ultra II Directional RNA Library Prep Kit for Illumina (NEB, E7760) according to manufacturer’s instructions.

### ChIP sequencing

The detailed ChIP-seq protocol was described (Kurachi et al., 2014). *Cross-linking the cells.* The in vitro cultured WT and Batf KO P14 CD8+ T cells were collected, and the cell numbers were counted. Cells were suspended in cell culture media with 1% formaldehyde and incubate the cells for 10 min at 37°C. Formaldehyde was quenched with glycine in a final concentration of 125 mM at room temperature for 5 min. The fixed cells were washed 2 times with ice cold PBS and the fixed cell pellets were freezed at −80 °C for storage. *Lyse the cells and shear the chromatin.* The frozen, fixed cells were thawed on ice for 10 min. Add 120 ul of fresh ChIP SDS Lysis Buffer (0.5% SDS, 50 mM Tris, pH 8, 10 mM EDTA, 1 × protease inhibitor cocktail (Roche)) per 5 million cells. Incubated the lysate on ice for 10 min and transferred to Covaris microTUBEs (Covaris, cat# 520045). The Covaris microTUBEs were transferred to the tube rack, placed into the Covaris E220 sonicator and sonicated the samples for 6 treatments (Peak power: 175.0, Duty factor: 10.0, Cycle/Burst: 200, Ave. Power:17.5) of 60 sec for each tube. *Prepare the IP beads.* Prepare the final ChIP buffer by adding 1-part ChIP SDS Lysis Buffer with 4 parts ChIP Dilution Buffer (1.25% Triton X-100, 12.5 mM Tris, pH 8, 187.5 mM NaCl) with protease inhibitors. 30-50 ul of Protein G Dynabeads (Invitrogen, cat# 10004D) were aliquoted into each ChIP tube, one for each ChIP up to 10 million cells. The beads were washed once in 300 ul of ChIP buffer and resuspended the beads in 500 ul of ChIP buffer. 2 to 10 ug of ChIP antibodies were added to the tube and the ChIP tubes were placed on a rotator at 4°C for 3 to 4 hours to bind the antibody to the beads. *Chromatin Immunoprecipitation.* After the samples have been sheared, pooled the lysates together in a microcentrifuge tube. Spin the pooled lysates at 13000 rpm, 4°C for 10 min. The cleared lysate was transferred to a new tube, added 4 parts of ChIP Dilution Buffer (with protease inhibitors) and mixed by pipetting up and down. 100 ul of lysate was aliquoted as an input fraction into another tube and stored in fridge at 4°C overnight. Once the antibodies were bound to the beads, quickly spun down the ChIP bead tubes (5-10 sec spin) and placed the tubes on a magnet rack. Aspirated the entire buffer and aliquoted an appropriate amount of ChIP lysate into the ChIP bead tubes. (Each 600 ul = 5 million cells). Placed the ChIP tubes on a rotator at 4°C overnight. *Prepare Elution Buffer.* Make fresh ChIP Elution Buffer: 1% SDS, 0.1 M sodium bicarbonate. *Washing and Elution of ChIP samples.* Removed the ChIP tubes from the rotator, quickly spun down the tubes, and placed the tubes on a magnet rack. Aspirated the lysate without disturbing the beads and then performed 5 washes for the ChIP beads with 1 ml of the following wash buffers for 3-5 min. Wash 1: ChIP Low Salt Buffer: 0.1% SDS, 1% Triton X-100, 20 mM Tris, pH 8, 2 mM EDTA, 150 mM NaCl. Wash 2: ChIP High Salt Buffer: 0.1% SDS, 1% Triton X-100, 20m M Tris, pH 8, 2 mM EDTA, 500 mM NaCl. Wash 3: ChIP LiCl Buffer: 0.7% sodium deoxycholate, 1% NP-40, 20m M Tris, pH 8, 1 mM EDTA, 500 mM LiCl. Wash 4: ChIP LiCl Buffer. Wash 5: ChIP TE Buffer: 10m M Tris, pH 8, 1 mM EDTA. After the wash steps, eluted the DNA twice with 100 ul ChIP Elution Buffer on a vortex mixer for 30 min. Tubes were removed from the vortex mixer, quickly spun down the tubes, and placed the tubes on a magnet rack. The samples were transferred to new tubes. Retrieved the input tubes from 4°C. Added 5 M NaCl to a final concentration of 0.2 M for each tube (input and ChIP samples). All tubes were placed (input and ChIP samples) on the heat block at 65°C overnight to reverse the formaldehyde cross-linking of the DNA and protein. *RNA/Protein Degradation* Removed samples from the heat block, added 2 ul of RNase A (Qiagen), and 2 ul of Proteinase K (RNA grade, Invitrogen/Life Technologies) to each tube. Tubes were placed on the heat block at 37°C for ~2 hrs. *Purify the ChIP DNA fragments.* Removed samples from the heat block, and added 10 ul of 3M sodium acetate, pH 5.5 to each input and ChIP sample. Qiagen MinElute Reaction Cleanup Kit was used to purify the DNA from the input and ChIP samples and eluted the DNA in 23 ul of EB buffer. Determine the concentration of ChIP DNA for library preparation using a Nanodrop. *ChIP-seq library construction.* NEBNext ChIP-Seq Library Prep Master Mix Set for Illumina (NEB cat# E6240L) was used for ChIP-seq library preparation. *End Repair of ChIP DNA*. Added up to 50 ng of ChIP DNA in a maximum of 20 ul to a PCR plate. Added 30 ul of the master mix (5 ul End Repair Reaction Buffer (10X), 24 ul nuclease free water, 1 ul End Repair Enzyme Mix) to each sample of ChIP DNA. Incubated the samples in a thermal cycler for 30 min at 20°C. Added 10 ul of 3M sodium acetate to each sample, purified the samples with Qiagen QIAquick PCR Purification Kit and eluted end-repaired DNA in 34 ul of EB. *dA-Tailing of End-Repaired DNA.* Added 16 ul of the master mix (5 ul dA-Tailing Reaction Buffer (10X), 10 ul nuclease free water, 1 ul Klenow Fragment) to each sample of 34 ul of end-repaired DNA. Incubated the samples in a thermal cycler for 30 min at 37°C. Added 10 ul of 3M sodium acetate to each sample. Use Qiagen MinElute Reaction Cleanup Kit to isolate and elute dA-Tailed DNA in 10 ul of EB. *Adaptor Ligation of dA-Tailed DNA and size-selection*. Diluted the NEBNext Adaptor (15 mM stock) 10X to a 1.5 mM working stock in nuclease free water. Added the 10 ul of dA-Tailed DNA to a PCR plate and added 20 ul of the master mix (6 ul Quick Ligation Reaction Buffer (5X), 1 ul Diluted Adaptor (1.5 mM), 9 ul nuclease free water, 4 ul End Repair Enzyme Mix) to each sample of ChIP DNA. Incubated the samples in a thermal cycler for 15 min at 20°C. Added 3 ul of USER enzyme to each sample and incubated the samples in a thermal cycler for 15 min at 37°C. After the incubation, used AMPure XP beads (Beckman Coulter A63881) for size selection of ChIP-seq DNA. Performed Double Size selection with beads to sample ratio: 0.5 for right side selection and 0.9 for left side selection. Eluted the DNA 20 ul of EB and the DNA is ready for amplification. *PCR Enrichment of Size Selected ChIP-Seq DNA.* NEBNext Multiplex Oligos for Illumina Index Primers Sets were used for indexing ChIP-seq libraries. Added the 2.5 ul of Universal PCR Primer and 2.5 ul index primers to samples. Added 25 ul of NEBNext HotStart Q5 2x PCR Master Mix to each sample and performed PCR with the following conditions: (1) 98°C 30 sec, (2)98°C 10 sec, (3) 98°C 10 sec, (4) 65°C 30 sec, (5) 72°C 30 sec (6) repeat (3)-(5) 12X, (7) 72°C 5 min, (8) hold at 4 °C. *SPRI Bead Clean Up of ChIP-Seq DNA.* Performed double size selection again with 0.55 for the right-side selection and 1.0 for the left-side selection. Eluted the samples with 40 ul Elution Buffer and examined the size range of your library using TapeStation. Antibodies used in ChIP-seq: Batf (Brookwood Biomedical, PAB4003), Jund (Santa Cruz, 329), Fosl2 (Santa Cruz, Q-20), Irf4 (Santa Cruz M-17), Stat3 (Santa Cruz, C-20), ETS-1 (Santa Cruz, C-20), T-bet (Santa Cruz, H-210), Stat5 (Santa Cruz A-9), Runx3 (Abcam, ab11905) and Eomes (Abcam, ab23345)

### ATAC-seq

50,000 cells were harvested and spun down at 500 g for 5 min, 4°C. The cell pellet was washed once with 50 μl of cold 1X PBS buffer and resuspend in 50 μl cold lysis buffer (10 mM Tris-HCl, pH 7.4, 10 mM NaCl, 3 mM MgCl2, 0.1% IGEPAL CA-630). Spun down immediately at 500 g for 10 min, 4°C. Supernatants were discarded, and proceeded to the transposition reaction. To make the transposition reaction mix, combined the following: 25 μl 2x TD Buffer (Illumina Cat #FC-121-1030), 2.5 μl Tn5 Transposase (Illumina Cat. #FC-121-1030), 22.5 μl nuclease free water to 50 μl of total volume. Resuspended nuclei in the transposition reaction mix and incubate at 37°C for 30 min. Immediately following incubation, purified the DNA using Qiagen MinElute Kit according to the manufacturer’s protocol and eluted transposed DNA in 20 μl Elution Buffer (10mM Tris buffer, pH 8). Purified DNA can be stored at −20°C. To amplify transposed DNA fragments and barcoded the libraries, Illumina Nextera Index Kit (FC-121-1011) was used. Combined the following in a PCR tube: 20 μl Transposed DNA, 2.5 μl Nextera PCR Primer with barcode 1, 2.5 μl Nextera PCR Primer with barcode 2, 25 μl NEBNext High-Fidelity 2x PCR Master Mix (New England Labs Cat. #M0541) to 50 μl total volume. PCR cycle as follows: (1) 72°C for 5 min, (2) 98°C 30 sec, (3) 98°C 10 sec, (4) 63°C 30 sec, (5) 72°C, 1 min, (6) Repeat steps 3-5, 12x (7) Hold at 4°C. Used AMPure XP Beads to purify the ATAC-seq libraries with a 1:1 PCR product to Beads ratio and followed manufacturer’s instructions. Eluted the purified ATAC-seq libraries in 40 μl Elution Buffer.

### Library quantification and sequencing

KAPA Library Quantification Kit for Illumina Platforms (KAPA Biosystems, KK4824) was used for sequencing library quantification according to the manufacturer’s protocol with ABI ViiA 7 Real-Time PCR System. NextSeq 500 V2 High Output Kit (Illumina, FC-404-2005, 75 cycles) and Illumina NextSeq 550 system were used for library sequencing.

### HiChIP

The detailed HiChIP protocol was described (Mumbach et al., 2016).

*Cell crosslinking* Resuspend every 1 million target cells in 1 ml 1% formaldehyde and incubate cells at room temperature for 10 minutes with rotation. Add glycine (125 mM final) to quench formaldehyde, and then incubate at room temperature for 5 minutes with rotation. Wash the cells once in PBS and then store in −80°C or proceed into the HiChIP protocol. *Lysis and restriction digest*. Resuspend 20 million crosslinked cells in 2 tubes of 500 μl of ice-cold Hi-C Lysis Buffer (10 mM Tris-HCl pH 7.5, 10 mM NaCl, 0.2% NP-40 and protease inhibitors) and rotate at 4 °C for 30 minutes. Spin down at 2500 g for 5 minutes at 4 °C and discard the supernatant. Wash pelleted nuclei once with 500 μl of ice-cold Hi-C Lysis Buffer. Remove the supernatant and resuspend the pellet in 100 μl of 0.5% SDS. Incubate at 62 °C for 10 minutes and add 285 μl of H2O and 50 μl of 10% Triton X-100 to quench the SDS. Mix well and incubate at 37 °C for 15 minutes with rotation. Add 50 μl of 10X NEB Buffer 2 and 375 U (15 μl of 25 U/ μl) of MboI restriction enzyme (NEB, R0147), and digest chromatin for 2 hours at 37°C with rotation. Heat inactivated MboI at 62 °C for 20 minutes. *Incorporation and proximity ligation.* To fill in the restriction fragment overhangs and mark the DNA ends with biotin, add 52 μl of fill-in master mix: 0.4 mM biotin-dATP (Thermo 19524016) 37.5 μl, 10 mM dCTP 1.5 μl, 10 mM dGTP 1.5 μl, 10 mM dTTP 1.5 μl, 5U/ μl DNA Polymerase I, Large (Klenow) Fragment (NEB, M0210) 10 μl. Mix and incubate at 37 °C for 1 hour with rotation. Add 948 μl of ligation master mix: 150 μl 10X NEB T4 DNA ligase buffer (NEB, B0202), 125 μl 10% Triton X-100, 3 μl 50 mg/mL BSA (Thermo Fisher, AM2616), 10 μl 400 U/ μl T4 DNA Ligase (NEB, M0202), 660 μl nuclease free water. Incubate at room temperature for 4 hours with rotation. Pellet nuclei at 2500 g for 5 minutes and remove supernatant. *Sonication.* For Batf and Ctcf HiChIP, pool the two sample tubes into 880 μl Nuclear Lysis Buffer and proceed to the sonication step. Transfer to a Covaris milliTUBE (Covaris #520135) and shear with Covaris E220 with the following program: Fill Level 5, Duty Cycle 5, PIP 140, Cycles/Burst 200, Time 4 minutes. Preclearing, immunoprecipitation, IP bead capture, and washes. Centrifuge sample for 15 minutes at 16000 g at 4°C. Add 3320 μl Dilution Buffer (0.01% SDS, 1.1% Triton X-100, 1.2 mM EDTA, 16.7 mM Tris pH 7.5, 167 mM NaCl) to the 880 μl sonicated sample and split into 4 tubes, 1050 μl per tube (~5 million cells). Wash 30 μl of Protein G Dynabeads (Invitrogen, 10004D) for every 5 million cells in 50 μl ChIP Dilution Buffer. Resuspend Protein G Dynabeads in 50 μl of Dilution Buffer per tube (100 μl per HiChIP), add to sample and rotate at 4 °C for 1 hour. Put samples on magnet and transfer supernatant into new tubes. Add 2.5 μg of antibody for every 5 million cells and incubate at 4°C overnight with rotation. Wash 30 μl of Protein G Dynabeads for every 5 million cells in ChIP Dilution Buffer. Resuspend Protein G Dynabeads in 50 μl of Dilution Buffer (100 μl per HiChIP), add to sample and rotate at 4 for 2 hours. Combine all the split samples and wash beads four times each with Low Salt Wash Buffer (0.1% SDS, 1% Triton X-100, 2 mM EDTA, 20 mM Tris-HCl pH 7.5, 150 mM NaCl), High Salt Wash Buffer (0.1% SDS, 1% Triton X-100, 2 mM EDTA, 20 mM Tris-HCl pH 7.5, 500 mM NaCl), and twice with the LiCl Wash Buffer (10 mM Tris-HCl pH 7.5, 250 mM LiCl, 1% NP-40, 1% Sodium deoxycholate, 1 mM EDTA). Washing was performed at room temperature on a magnet by adding 500 μl of a wash buffer, swishing the beads back and forth twice by moving the sample relative to the magnet, and then removing the supernatant. *ChIP DNA elution.* Resuspend ChIP sample beads in 100 μl of fresh DNA Elution Buffer (50 mM NaHCO3, 1% SDS). Incubate at room temperature for 10 minutes with rotation, followed by 3 minutes at 37 °C shaking. For ChIP samples, place on magnet and remove supernatant to a fresh tube. Add another 100 μl of DNA Elution Buffer to ChIP samples and repeat incubations. Remove ChIP samples supernatant again to the new tube. There should now be 200 μl of ChIP sample. Add 10 μl of Proteinase K and 8 μl of 5M NaCl to each sample and incubate at 55 °C for 45 minutes. Increase temperature to 67 and incubate for at least 1.5 hours with shaking. Use QIAGEN minElute Kit to purify the samples and elute in 12 μl of water. Quantify post-ChIP DNA with Quant-iT dsDNA HS Kit (ThermoFisher, Q33120). For libraries with greater than 150 ng of post-ChIP DNA, set aside material and take a maximum of 150 ng into the biotin capture step. *Biotin pull-down and preparation for Illumina sequencing.* Prepare for biotin pull-down by washing 5 μl of Streptavidin C-1 (Invitrogen, 65001) beads with Tween Wash Buffer (5 mM Tris-HCl pH 7.5, 0.5 mM EDTA, 1 M NaCl, 0.05% Tween-20). Resuspend the beads in 10 μl of 2X Biotin Binding Buffer (10 mM Tris-HCl pH 7.5, 1 mM EDTA, 2 M NaCl) and add to the samples. Incubate at room temperature for 15 minutes with rotation. Separate on a magnet and discard the supernatant. Wash the beads twice by adding 500 μl of Tween Wash Buffer and incubating at 55 °C for 2 minutes with shaking. Wash the beads in 100 μl of 1X TD Buffer (10 mM Tris-HCl pH 7.5, 5 mM MgCl2, 10% Dimethylformamide). Resuspend beads in 25 μl of 2X TD Buffer (20 mM Tris-HCl pH 7.5, 10 mM MgCl2, 20% dimethylformamide). For Batf and Ctcf HiChIP, use 2.5 μl Tn5 transposase for 50 ng post-ChIP DNA and add water to 50 μl. Incubate at 55 °C with interval shaking for 10 minutes. Place samples on magnet and remove supernatant. Add 50 mM EDTA to samples and incubate at 50 °C for 30 minutes, then quickly place on magnet and remove supernatant. Wash samples twice with 100 μl 50 mM EDTA at 50 °C for 3 minutes and wash samples twice in 100 μl Tween Wash Buffer at 55 °C for 2 minutes. And then wash samples in 100 μl 10 mM Tris. Resuspend beads in 20 μl Elution Buffer. PCR and Post-PCR Size Selection. Add 30 μl PCR master mix (2.5 μl Illumina dual indexes primer 1, 2.5 μl Illumina dual index primer 2, 25 μl NEB Q5 HotStart Master Mix) to the 20 μl beads bound with samples. For 50 ng post-ChIP DNA, Run PCR program: (1) 72°C 5 minutes, (2) 98°C 1 minute, (3) 98°C 15 seconds, (4) 63°C 30 seconds, (5) 72°C 1 minute, (6) repeat (3)-(5) 5X, (7) hold at 4°C. *Purify and size-select the PCR product with Ampure XP beads.* After PCR, place libraries on a magnet and elute into new tubes (~50 μl). Then add 25 μl of Ampure XP beads and keep the supernatant to capture fragments less than 700 bp. Transfer supernatant to a new tube and add 15 μl of fresh beads to capture fragments greater than 300 bp. Eluted in 22 μl Elution Buffer and the samples were ready to be quantified and sequencing. Antibodies used in HiChIP experiments: Batf (Brookwood Biomedical, PAB4003) and Ctcf (Millipore, 07729MI).

### Lentiviral vector construction and lentivirus production

The coding domain sequence (CDS) of mouse Batf, Irf4, Runx3 and T-bet were PCR amplified and cloned into pDONR221 vector to generate entry clones pENTR221-Batf, pENTR221-Irf4, pENTR221-Runx3 and pENTR221-T-bet respectively. The CDS of Batf, Irf4, Runx3 and T-bet from the entry clones were then cloned into the pLoxp404 lentiviral expression vectors (Broad Institute) pLoxP-404-HygroR, pLoxP-404-PuroR, pLoxP-404-BlastR and pLoxP-404-GFP by Gateway Cloning (Thermo Fisher Scientific) to generate final lentiviral constructs pLoxP-404-HygroR-Batf, pLoxP-404-PuroR-Irf4, pLoxP-404-BlastR-Runx3 and pLoxP-404-GFP-T-bet. The lentivirus was packaged in HEK 293T cells by co-transfecting the lentiviral vectors with pCMV-VSV-G (envelope) and pCMV-dR8.91 (gag-pol) plasmids. The supernatants containing lentivirus were harvested and used for NIH/3T3 cell line transduction. The GFP-T-bet lentivirus was concentrated with ultracentrifugation and resuspended in medium as 50X concentrated virus for transduction.

### Lentiviral transduction, antibiotics selection and doxycycline induction of NIH/3T3 cells

24 hours before transduction, seed NIH/3T3 cells in 6-well plates, 2 × 10^5^ cells/well in 2 ml DMEM (10% FBS) medium. On the day of transduction, replace with fresh medium and add 300 ul lentiviral supernatant with 5 ug/ml polybrene. After overnight incubation, replace the virus containing medium with fresh DMEM and incubated for an additional 48 hours before adding antibiotics. 200 ug/ml Hygromycin B, 10 ug/ml Blasticidin, 7.5 ug/ml Puromycin were added to corresponding transduced 3T3 cells and selection for 1 to 2 weeks for generating stable clones. The transduction efficiency of GFP-T-bet lentivirus transduced 3T3 cells was determined by flow cytometry for GFP expression and GFP expressing cells were sorted if necessary. 4 ug/ml doxycycline was used to induce the gene expressions in transduced 3T3 fibroblasts. The cells were treated for 72 hours and the culture medium was changed every 24 hours with fresh doxycycline.

### Western blot

Fibroblasts were trypsinized, washed with cold PBS and lysed in RIPA lysis buffer (Thermo Scientific) on ice for 30 min for extracting protein lysates. Samples were centrifuged >13,000 rcf at 4 °C for 10 min, supernatants were collected and protein concentrations were quantified with Bio-Rad protein assay (Bio-Rad). 20-40 ug of lysates were mixed with LDS sample buffer (ThermoFisher) and heated at 70 degrees for protein denaturation according to the manufacturer’s protocol. Samples were run in Bolt 4-12% Bis-Tris Plus Gels (ThermoFisher) and transferred to PVDF membrane (Millipore). The following primary antibodies were used to detect designated proteins: Batf (Brookwood Biomedical, PAB4003), Irf4 (Santa Cruz, M-17), Runx3 (Abcam, ab11905), T-bet (Santa Cruz, H-210), TGFb1 (Abcam, EPR18163) and Hsp90 (Santa Cruz, F-8). HRP conjugated secondary antibodies (Goat anti-rabbit IgG and Goat anti-mouse IgG light chain) were purchased from Jackson ImmunoResearch Laboratories. HRP substrates (Immobilon western chemiluminescent HRP substrate, Millipore) were applied to the membrane according to manufacturer’s protocol and imaged using Fujifilm LAS4000 imaging system.

### Flow cytometry

Fibroblasts were trypsinized (Trypsin-EDTA 0.25%, Gibco), washed with cold PBS and resuspended in flow cytometry staining buffer (eBioscience). The cells were stained at 4 degrees for 30 mins with the following fluorochrome-conjugated antibodies: APC-IL17Ra (eBioscience, clone PAJ-17R) and APC-CD197(CCR7) (BioLegend, clone 4B12). Samples were washed with flow cytometry staining buffer and analyzed on LSRFortessa (BD). The data were analyzed with FlowJo software (FlowJo).

### Alignment of RNA-seq and quantification of transcript abundance

All RNA-Seq reads were trimmed using Trimmomatic (Bolger et al., 2014) to remove primer and low-quality bases. Reads were then passed to FastQC [http://www.bioinformatics.babraham.ac.uk/projects/fastqc/] to check the quality of the trimmed reads. The single-end reads were then aligned to the Mus_musculus.GRCm38.82 transcriptome from Ensembl using RSEM (Li and Dewey, 2011) with the parameters “--num-threads 4 --bowtie2 --sampling-for-bam --output-genome-bam --sort-bam-by-coordinate --sort-bam-memory-per-thread 1G --estimate-rspd --fragment-length-mean 200 --fragment-length-sd 80’.

### Analysis of differential gene expression

Differential expressions were assessed using DESeq2 (Love et al., 2014). Genes were considered differentially expressed if they had an adjusted p-value < 0.05. ‘betaPrior’ was set equal to FALSE.

### Analysis of time course in vitro RNA-Seq

Using DESeq2 (Love et al., 2014), we conducted a likelihood ratio test to identify any genes that changed over time (adjusted p value <0.05). We then clustered the genes using the pheatmap package in R with correlation as the row distance, and hierarchical complete clustering (default settings), cutting the tree at the point at which there were five clusters.

### Alignment of ATAC-seq and peak calling

All ATAC-seq reads were trimmed using Trimmomatic (Bolger et al., 2014) to remove primer and low-quality bases. Reads were then passed to FastQC (Andrews, 2010) to check the quality of the trimmed reads. The paired-end reads were then aligned to the mm10 reference genome using bowtie2 (Langmead and Salzberg, 2012), allowing maximum insert sizes of 2000 bp, with the “--no-mixed” and “--no-discordant” parameters added. Reads with a mapping quality (MAPQ) below 30 were removed. Duplicates were removed with PicardTools, and the reads mapping to the blacklist regions and mitochondrial DNA were also removed. Reads mapping to the positive strand were moved +4 bp, and reads mapping to the negative strand were moved −5bp following the procedure outlined by Buenrostro et al. (Buenrostro et al., 2013) to account for the binding of the *Tn5* transposase.

Peaks were called using macs2 on the aligned fragments (Zhang et al., 2008) with a q-value cutoff of 0.05 and overlapping peaks among replicates were merged. A region was considered a valid peak if it had a q-value below 0.05 in at least two replicates, and a qvalue below 0.001 in at least one of the replicates.

### Tests of differential accessibility in ChARs

Differential accessibility was assessed by applying DESeq2 (Love et al., 2014) to a matrix with rows as peaks, columns as samples, and counts of Tn5 cuts in each cell. Peaks were considered differentially accessible if they had an adjusted p-value < 0.05. ‘betaPrior’ was set equal to FALSE.

### Alignment of ChIP-seq and peak calling

All ChIP-seq reads were trimmed using Trimmomatic (Bolger et al., 2014) to remove primer and low-quality bases. Reads were then passed to FastQC (Andrews, 2010) to check the quality of the trimmed reads. These single-end reads were then aligned to the mm10 reference genome using bowtie2 (Langmead and Salzberg, 2012), allowing maximum insert size of 2000 bp, with the “--no-mixed” and “--no-discordant” parameters added. Reads with a mapping quality (MAPQ) below 30 were removed. Duplicates were removed with Picard Tools, and the reads mapping to the blacklist regions and mitochondrial DNA were also removed.

Each combination of transcription factor and condition had two replicates, with the exception of Runx3-WT, which we limited to one high-quality replicate. ChIP-Seq peaks were called in each replicate, versus a control sample, using macs2 (Zhang et al., 2008) with a q-value cutoff of 0.05 and overlapping peaks among replicates were merged. A region was considered a valid peak for a transcription factor if it had a q-value below 0.05 in at least two replicates, and a qvalue below 0.001 in at least one of the replicates.

### Tests of differential binding

Differential binding was assessed using the csaw package (Lun and Smyth, 2016) in regions identified as being bound in either the WT or *Batf^−/−^* for the given transcription factor. Only log2 fold changes greater than one with an adjusted p-value below 0.05 were considered differentially bound.

### Universe of transcription factor bound regions and accessible peaks

The universe of transcription factor bound regions and accessible peaks was created by merging the overlapping ChIP-seq and ATAC-seq peaks previously described. ATAC-seq peaks from the experiments in vivo flox experiments were used, and All WT and Batf KO ChIP-seq peaks previously described were used. This dataset is available as <u>Table 2</u>.

### Test for Association of transcription factor binding with reduced accessibility of ChARs in Batf cKO

We used the 51,259 regions from Table 2 where an ATAC-seq peak was present in the WT (floxed) or Batf cKO condition. We used the results from the test of differential accessibility (described above) to code each peak as having reduced accessibility in Batf cKO (1) or not (0). We then ran a separate logistic regression for each of our TF’s (coded 1 if bound, 0 if not bound) to test for its association with reduced accessibility in Batf cKO. All TF’s had a Bonferroni corrected p-value <0.001.

### Fold enrichment of transcription factor binding

Fold-enrichment of individual transcription factors is shown on the diagonal (barplot) of Figure 2F, and fold-enrichment for co-binding of transcription factors is shown on off-diagonal cells. X = locus bound by column transcription factor, Y = locus bound by row transcription factor, and L = locus less accessible in Batf KO. Fold-enrichment is calculated as:

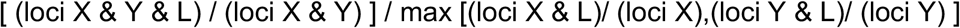

### Alignment of HiChIP data and peak calling

HiChIP data were aligned using the HiCPro pipeline (Servant et al., 2015). Normalized counts were plotted using the HiCPLotter python package (Akdemir and Chin, 2015).

The Hichipper package (Lareau and Aryee, 2018) was used to identify loops in the HiChIP data. The Hichipper package uses the Mango package (Phanstiel et al., 2015) to test the statistical significance of loops. We retained loops with an adjusted p-value <0.001 and at least 5 paired-end tags linking the two anchors.

### Gene-level enrichment analysis

Gene Set Enrichment Analysis was conducted with clusterProfiler (Yu et al., 2012) which utilizes fgsea (Sergushichev, 2016). The gsea, gseGO, and gseKEGG functions were run with 1000 permutations and an adjusted p-value cutoff of 0.10. Results are provided for the ImmSigDB C2 and C7 collections (min set size = 10, max set size = 1000), GO Biological Processes and Molecular Functions (min set size = 100, max set size = 500), and KEGG (min set size = 100, max set size = 500). The “simplify” function was run on the GO results to condense them.

The Wald statistic from DESeq2 was used to order the genes. Mouse ensembl gene id’s were mapped to human orthologs using Biomart (Durinck et al., 2005, 2009) for the C2 and C7 collections. For instances in which multiple mouse genes mapped to one human ortholog, the average of the Wald statistics was used. Genes which were knocked out or ectopically expressed were removed prior to GSEA to avoid biasing the results.

### T cell score calculation and Shapley value

T cell gene expression signatures were obtained from DESeq2 comparison of RNA-seq of effector and naive CD8+ T cells vs fibroblasts. Signature was defined by giving a weight of +1 to the top 1000 genes differentially upregulated in T cells and weight of −1 to the top 1000 genes differentially upregulated in fibroblasts. Weights were then applied to the vector of normalized counts for each condition. Shapley value for each TF was calculated based on the relative T cell score. Bootstrap estimates were calculated by computing the score using one randomly chosen replicate from each condition, with 2,500 iterations.

### Locus- level enrichment analysis

Subsets of loci deemed “foregrounds” were analyzed for fold enrichment of features relative to a background set of loci. The features analyzed were motifs and other genomic annotations, SNPs associated with immune diseases, proximity to gene groups of interest, and pathways in GREAT. q-values were estimated using the q-value package (Storey JD, Bass AJ, Dabney A, Robinson D, 2019).

### Motifs / annotation tracks

PWM’s for motifs were downloaded from the 2018 release of JASPAR (Khan et al., 2017). We used fimo (Grant et al., 2011) to identify motifs in mm10, and applied the default threshold of 1e-4. We also included regulatory features from the ORegAnno database (Lesurf et al., 2016), (iii) conserved regions annotated by the multiz30way (Blanchette et al., 2004) algorithm, and repeat regions annotated by RepeatMasker (Smit et al., 1996).

### SNPs

SNPs associated with different immune diseases were retrieved from the experimental factor ontology (Malone et al., 2010) by downloading all SNPs in the GWAS Catalog (Buniello et al., 2019) corresponding to a given node and its children. The orthologous regions in mice were identified by using liftOver (Hinrichs et al., 2006).

### GREAT pathways / genes

Loci were associated with pathways using GREAT (McLean et al., 2010), submitted with the rGREAT package (Gu, 2015). We retrieved pathways found in the MSigDB Immunologic Signatures, MSigDB Pathway, and GO Biological Process databases. Loci were mapped to genes using GREAT.

## ACKNOWLEDGMENTS

We thank Dr. Robert Manguso and Kathleen Yates of the Haining laboratory and members of Yosef laboratory for comments and suggestions, the Birgit Knoechel laboratory of Dana-Farber Cancer Institute for sequencing assistance, and the Hematologic Neoplasia and Jimmy Fund Flow Cytometry Cores of Dana-Farber Cancer Institute for technical support. This work was supported by National Institutes of Health (R01AI115712 to H.W.N. and W.E.J.), and the “Postdoctoral Research Abroad Program” of the Ministry of Science and Technology, Taiwan (MOST 104-2917-I-564-018 to H.W.T.)

## Author Contributions

H.W.T., J.K., W.N.H. and N.Y. designed the research and wrote the manuscript.

H.W.T. performed most of the experiments.

J.K. performed most of the computational analyses.

M.K. conducted the in vivo LCMV challenge and sorted the cells for this study.

R.A.B., M.A.D. and M.W.L. assisted the ChIP-seq and ectopic TF expression experiments.

W.I. and T.K. provided the critical mouse model for this study.

N.Y., W.N.H. and E.J.W.supervised the study.

