## Supplemental Figures for "Batf-mediated Epigenetic Control of Effector CD8+ T Cell Differentiation"

#### Supplementary Figure 1

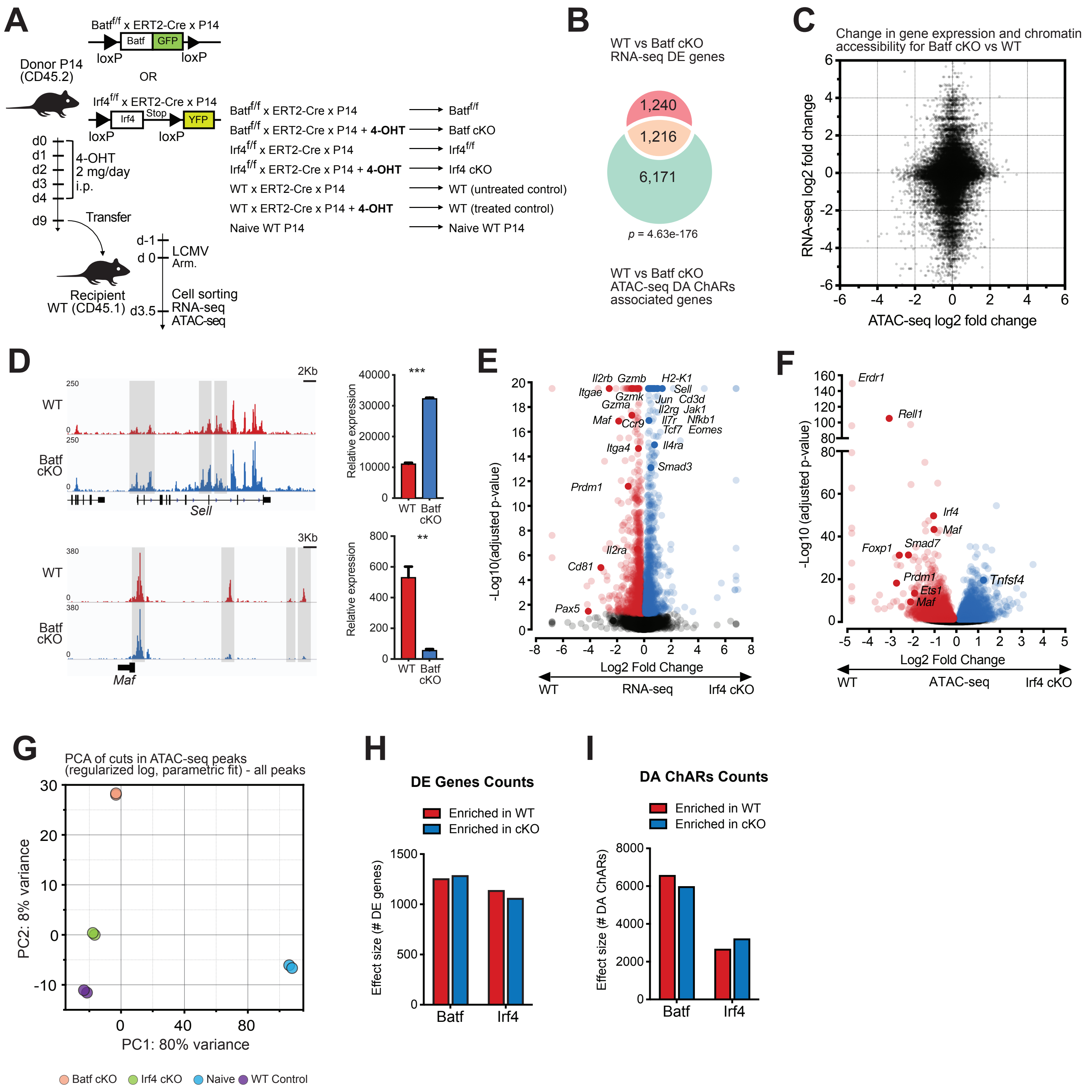

#### Supplementary Figure 1 (continued)

J

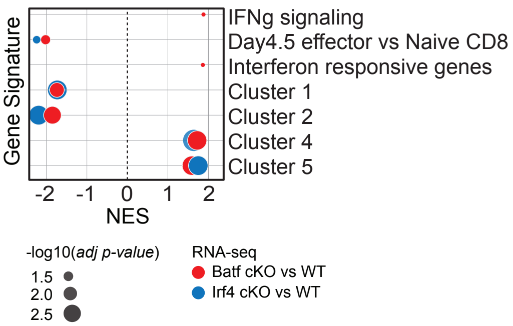


K

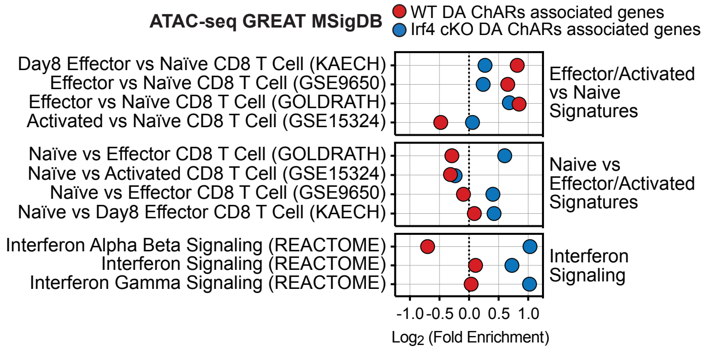


L

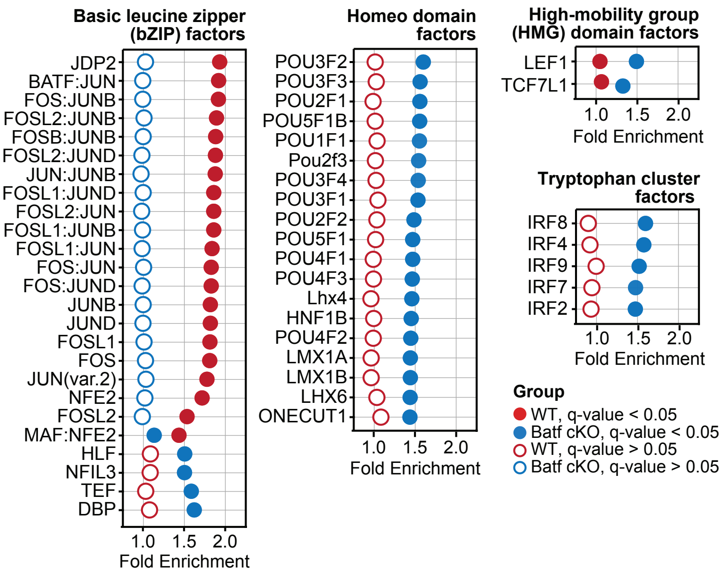

#### Supplementary Figure 2

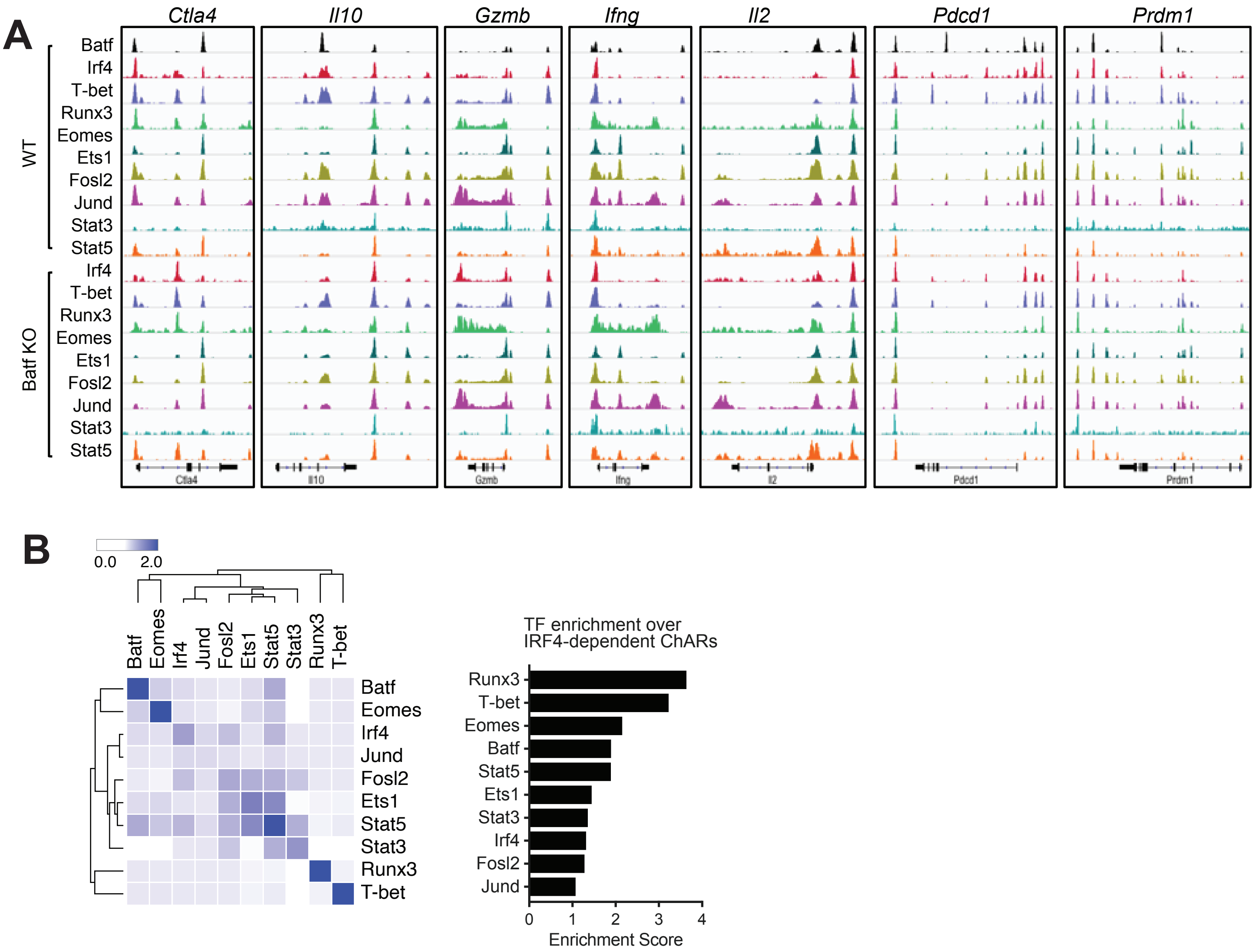

### Supplementary Figure 3

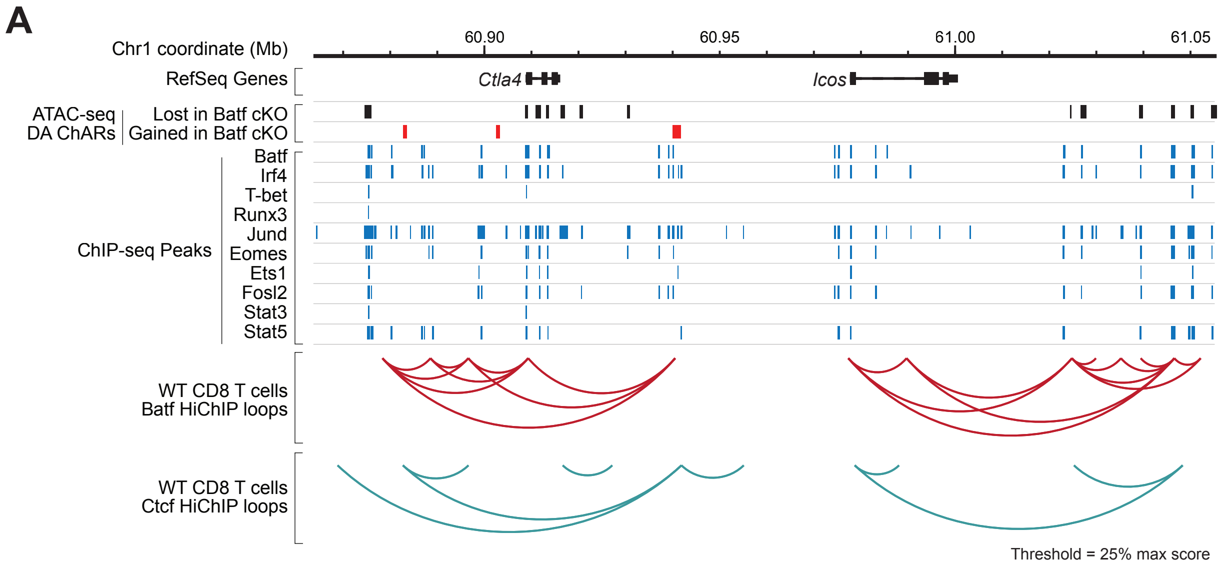

**Supplementary Figure 4**

A

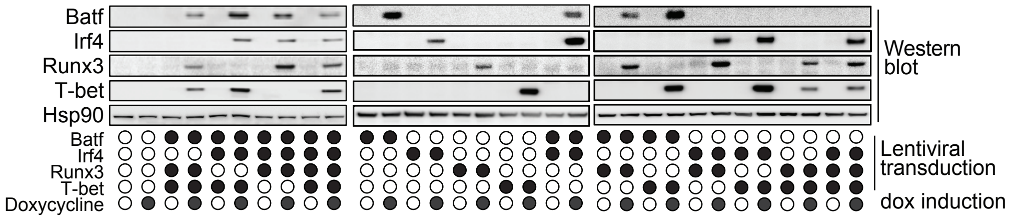


B

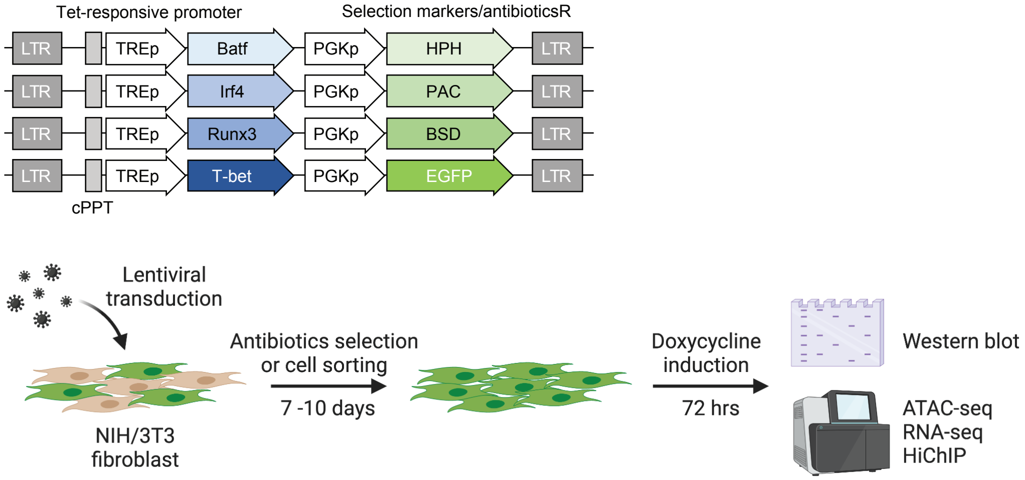


C

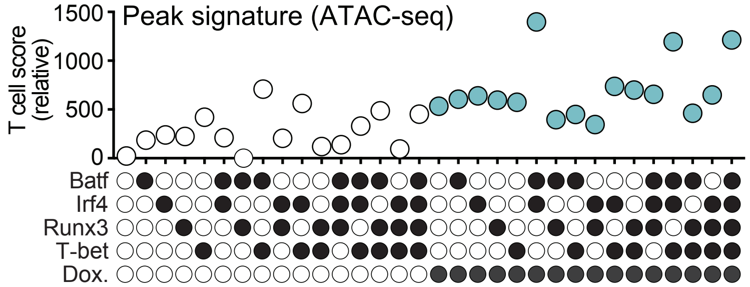

### Supplementary Figure 5

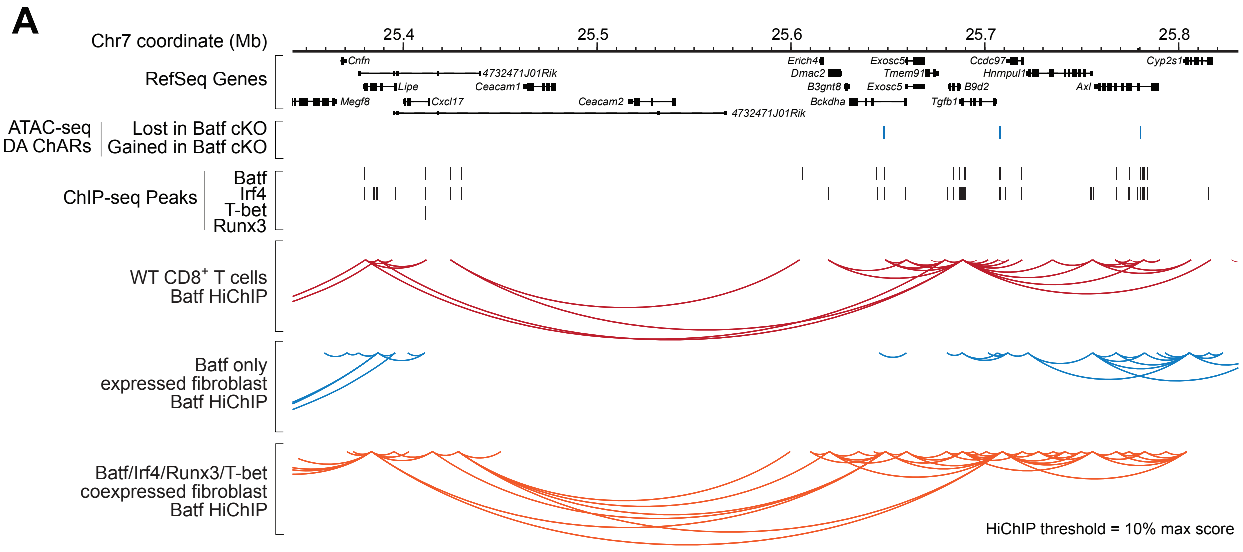


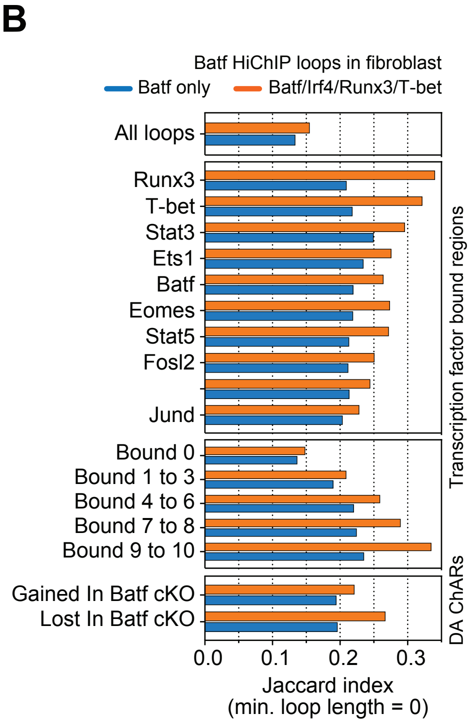

### Supplementary Figure 6

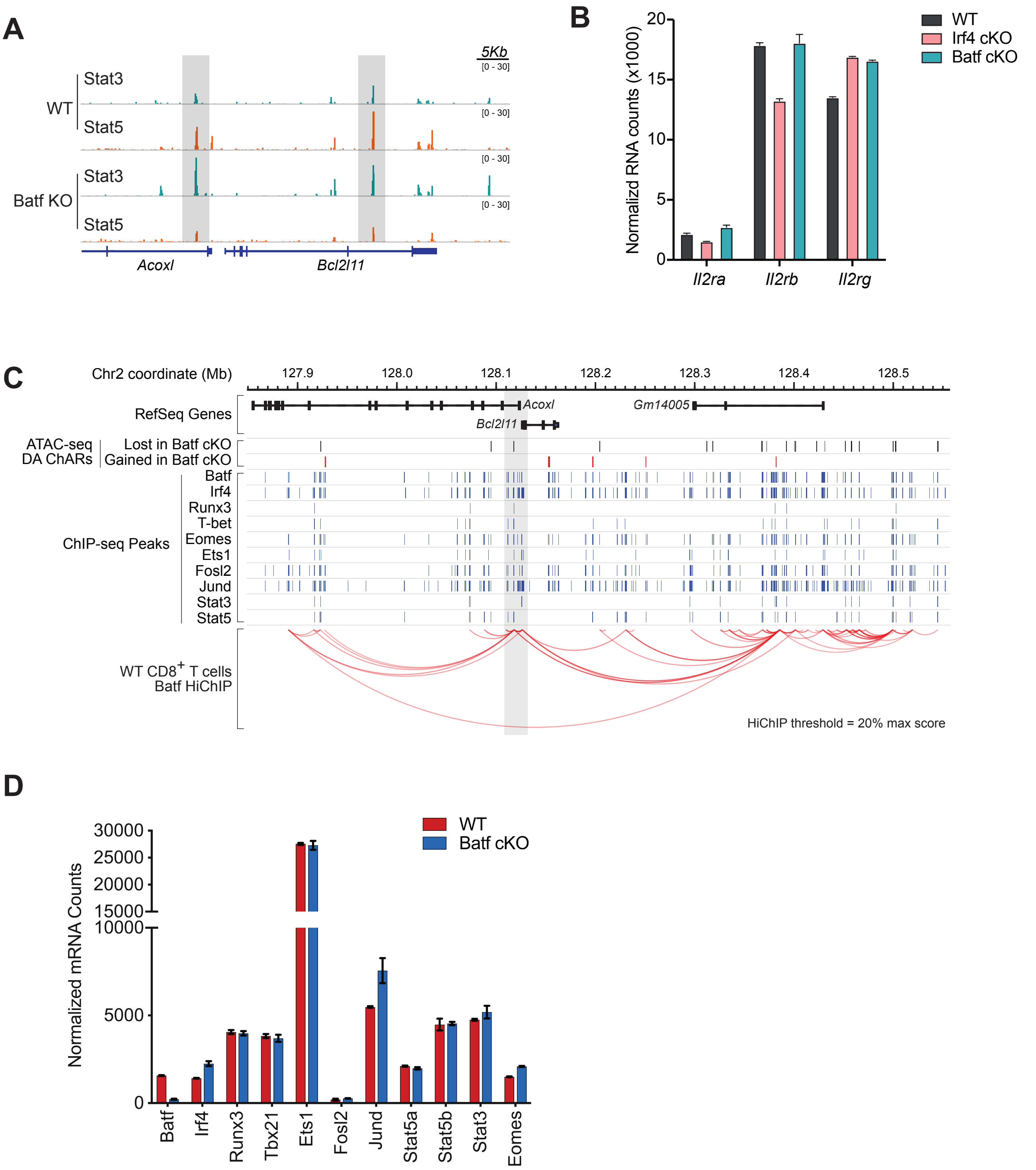
